# Structural changes in autism reflect atypical brain network organization and phenotypical heterogeneity: a deep network approach

**DOI:** 10.1101/2025.11.02.686152

**Authors:** Siddharth Lokray, Basilis Zikopoulos, Arash Yazdanbakhsh

## Abstract

Autism Spectrum Disorder (ASD) is a developmental disorder characterized by heterogeneity in social and emotional responses, language, and behavior. Assessments such as the Social Responsiveness Scale (SRS) can quantify this variability but understanding underlying mechanisms and identifying distinct and shared atypical organization and function of brain networks remains a challenge. Convolutional Neural Networks (CNNs) have been used to analyze imaging data. However, the relationship between structural brain changes observed in structural MRI (sMRI), the affected brain functional networks inferred from these structural changes, and their connection to ASD phenotypes and scores still requires systematic investigation. In this study, we used 3D CNNs to conduct a comprehensive analysis of macrostructural changes in ASD. We found consistently dominant involvement of (a) the left hemisphere, (b) the frontal and temporal lobe, and (c) the default mode network and dorsal/ventral attention networks in ASD. Our systematic AI approach utilized the phenotypic heterogeneity and spectral nature of ASD to uncover significant structural changes across brain regions and functional networks, correlating the structural, functional, and phenotypical heterogeneity of individuals with ASD. This enabled us to identify known and novel global and local brain region and network changes in ASD in relation to phenotypes and clinical scores that can guide diagnostic subtyping.

**Teaser (one sentence summary):** We used machine learning approaches to characterize the phenotypic and imaging-derived structural heterogeneity of ASD with the aim to identify the brain regions and networks linked to its cognitive and behavioral phenotypes.

## Introduction

Autism Spectrum Disorder (ASD) is a developmental disorder that affects social interactions, emotional responses, language, and speech development, leading to behavioral inflexibility in affected individuals (Amaral, Schumann, and Nordahl 2008; Ecker 2012; Lord et al. 2000). Types, onset, and severity of symptoms can vary significantly; thus, ASD affects individuals differently leading to a display of a variety of phenotypical combinations. This heterogeneity makes it challenging to understand the underlying mechanisms of disruption, pinpoint cortical and subcortical regions that comprise atypical brain networks in ASD, and ultimately accurately diagnose and treat affected individuals. Heterogeneity in ASD can be partially expressed as a function of the cognitive, behavioral, and verbal scores in the Autism Diagnostic Interview (ADI) or Social Responsiveness Scale (SRS) (Ecker 2012; Lord, Rutter, and Le Couteur 1994). A key question is whether ASD heterogeneity indices, based on these scores, can be reliably and consistently reflected in and correlated with the structural and functional changes in brain networks that can be assessed *in vivo* by imaging techniques, such as Magnetic Resonance Imaging (MRI) (Amaral, Andrews, and Nordahl 2024; Duan et al. 2025) .

Modern AI techniques, including 2- and 3-Dimensional versions of Convolutional Neural Networks (CNNs) and their derivatives, deep attentional, contrastive, and CNN and Long short-term memory (LSTM) hybrids have been used to analyze imaging data (Aghdam, Sharifi, and Pedram 2019; Aglinskas, Hartshorne, and Anzellotti 2022; Ecker et al. 2010; Khosla et al. 2018; Li et al. 2018; Niu et al. 2020; Supekar et al. 2013; Uddin et al. 2013; Zhao et al. 2018). Most of these studies focused on accurately classifying ASD against typically developed (TD) individuals, using structural imaging datasets, or identified disruptions in functional brain network activity predominantly using functional imaging (fMRI). For example, Aglinskas et al. (Aglinskas et al. 2022) developed a neural network that identified ASD- and control-specific neuroanatomical variability and correlated ASD anatomical features with symptoms. Using resting state fMRI data, Zhao et al. (2018) developed a neural network to classify between ASD and TD subjects and identified brain regions with the highest contribution to classification that were then associated with functional brain networks that are disrupted in ASD. However, two key relationships remain poorly explored and understood: a) the presence of structural changes and their correlation across brain areas, as a structural pointer to the functionally affected brain networks, and b) correlations between autism diagnostic scores and brain pathology (Uddin et al. 2013).

To address these gaps, we investigated the relationship between structural changes across regions and SRS Total scores and their association with functional brain networks to reveal common and divergent structural and functional changes in individuals with high- and low-severity ASD. Our overarching hypothesis is that sMRI, in fact, contains information that reflects functional network and diagnostic score changes in ASD, which can be deciphered, using state-of-the-art AI, to identify relevant brain regions involved in ASD severity prediction and their functional networks underlying ASD heterogeneity. This approach aims to demonstrate that network level markers of ASD pathology, which conventionally require functional imaging methods, can also be recovered from structural MRI. Moreover, our approach aims to extend our current understanding of ASD by linking specific brain regions to specific symptom scores and characterizing ASD severity as a continuum (high-/low-severity). Our findings suggest that there is indeed partial required information available in macroscopic sMRI that can be used to reliably identify brain regions, covariation, and functional networks involved across the heterogeneous spectrum of ASD, and provides a framework for the identification of systematic, correlated neuroanatomical and functional dimensions across ASD.

## Results

In this study, we examined sMRI datasets (ABIDE II, Table S1) from individuals with different ages and different ASD severity scores to capture ASD heterogeneity. Our approach involved development and optimization of CNNs to evaluate entire brain volumes, allowing for the 3D CNN to extract features to distinguish between high- and low-severity ASD individuals based on their structural MRIs. To reduce possible confounds, we used leave-one-site-out cross-validation, confound screening (age/eTIV/IQ/Euler/site), and label-permutation test. We then extracted brain regions with the highest importance for ASD heterogeneity utilizing Integrated Gradients (Sundararajan et al. 2017) and projected the extracted gradient measures on cortical parcellations using the Destrieux atlas (Destrieux et al. 2010). This enabled a more detailed identification of brain regions with macro-structural changes as highlighted by gradient-based saliency maps, along with the analysis of correlations between these changes, and the severity of ASD. Saliency seed stability, morphometric comparisons, and model randomization test were used to confirm that extracted saliency maps from the 3D CNN were reproducible and meaningful. We then identified hierarchical relationships between brain regions, based on our correlation analysis, to associate correlated regions with functional brain networks. These relationships were supported by inter-structural-covariance-analysis (ISCA) against morphometric covariance. We focused on functional networks that are affected in ASD, including the default mode network (DMN), salience network (SN), social network, and central executive network (CEN or frontoparietal network), which play key roles in self-awareness, attention, social interactions, and cognitive control. We also included the Language, Sensorimotor, Dorsal/Ventral Attention, and Limbic/Emotion networks, known to be affected in ASD, in the same analysis.

We used a pretrained MedicalNet (3D ResNet 50) backbone (He et al. 2015, Chen et al. 2019) with frozen weights and a trainable classification head on T1-weighted sMRI scans from individuals with high- and low-severity ASD (Fig. 1). The classification head was trained using the Adam optimizer, with a learning rate of 3×10^-5^, and batch size of 4. We screened for confounds including age, estimated total intracranial volume (eTIV), IQ, site, and Euler number (Table S1-S3). Age and eTIV did not differ between high- and low-severity groups (age: p_FDR = 0.173; eTIV: p_FDR = 0.524). However, IQ and Euler number did differ between groups after FDR correction (IQ: p_FDR = 8.1×10^-5^, Euler number: p_FDR = 8.9×10^-5^). Site distribution was nominally associated with severity group (χ^2^ = 9.82, p = 0.044) but did not survive FDR correction (p_FDR = 0.073). IQ differences are considered further in the Discussion, as there is a known association between IQ and ASD symptom severity. Given the site imbalance, we used a five-fold leave-one-site-out cross-validation (site abbreviations: KKI, NYU, OHSU, SDSU, TCD) with adaptive batch normalization (AdaBN) to address site specific differences as well as uneven high-/low-severity distributions per site (see Table S1, S3).

**Figure 1:**
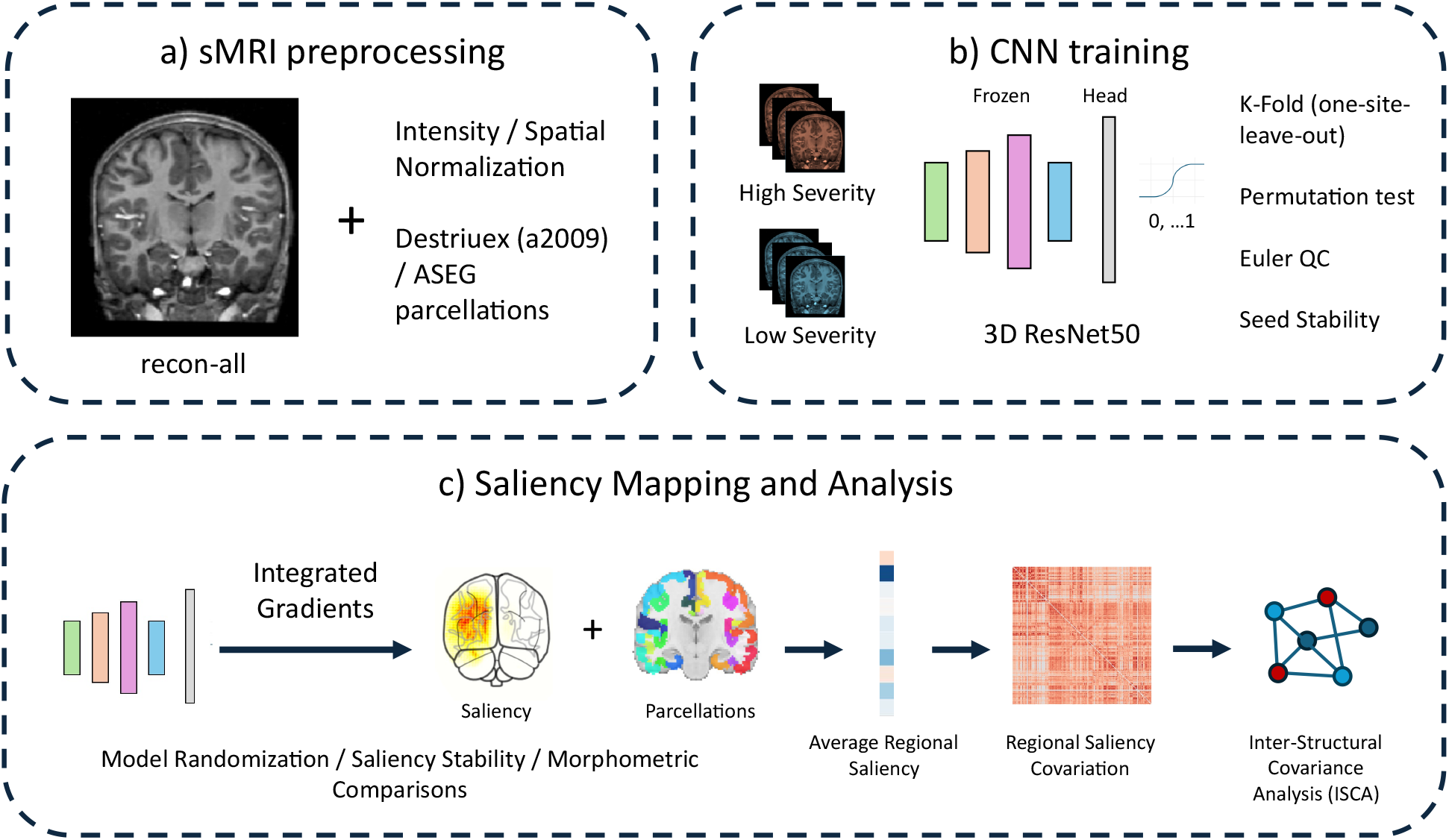
Overview of structural MRI preprocessing, model training, and saliency-based network analysis used to correlate ASD symptom severity with structural MRI. a) sMRI preprocessing: T1-weighted sMRI were processed through FreeSurfer’s recon-all pipeline for intensity and spatial normalization as well as cortical (Destrieux, a2009s) and subcortical (ASEG) parcellation. b) CNN training: preprocessed high- and low-severity ASD cases, defined by SRS (Social Responsiveness Scale) total score, were used to train a MedicalNet (3D ResNet-50) with a frozen backbone and a trainable classification head to predict ASD severity. Model validity was assessed by leave-one-site-out k-fold cross-validation, a label-permutation test, Euler-number seed stability, and seed-stability analysis across random initializations and data splits. c) Saliency mapping and analysis: per-subject Integrated Gradients attribution maps were computed using each fold’s own held-out model and validated with model-randomization, saliency-stability, and morphometric-comparison sanity checks. Saliency maps were co-registered with atlas parcellations from a) to compute each subject’s average regional saliency. Pairwise Kendall’s τ correlations between regions’ average saliency across subjects were then used to build a regional saliency covariation matrix, supported by ISCA, and visualized as a force-directed (spring-cluster) network graph to identify significant network involvement.

Over 1000 seed stability runs (varying data split and classification-head initialization), the 3D CNN achieved a mean balanced accuracy of 0.609 ± 0.029, consistent with an independent label-permutation test (1000 permutations: null mean = 0.500 ± 0.056; empirical p = 0.030). Given the significant Euler-number difference between low- and high-severity groups, we tested whether scan-quality variation could account for classification performance. The 3D CNN’s output was not correlated with Euler number (r = -0.027, n.s.), and Euler number could not be decoded from the frozen backbone’s features above chance across 1000 seed/split runs (balanced accuracy = 0.509 ± 0.029). Together these results indicate that Euler-number-related scan-quality variation is not encoded in the 3D CNN, and thus, cannot be contributing to the CNN’s severity classification.

Gradient-based saliency maps for downstream analysis were extracted using the trained classification-head with the lowest validation loss for each leave-one-site-out fold (Table S4). Across five folds, this selection criterion yielded a mean balanced accuracy of 0.636. To verify saliency maps reflected learned task-relevant weights, we compared regional saliency from the trained model against saliency generated after fully randomizing the network’s weights. Full randomization appropriately collapsed the relationship to the trained model’s saliency at the regional level (mean signed cosine similarity = -0.197, p = 0.155; mean unsigned cosine similarity = 0.449, p = 0.103), consistent with saliency reflecting task-relevant structures. Saliency map reproducibility was assessed by computing regional saliency for 10 subjects across 100 seed/split pairs each. Regional saliency maps were highly consistent across seeds (mean signed cosine similarity = 0.98, p = 0.001), indicating that gradient-based saliency was reproducible and not an artifact of random initialization.

We used the Destrieux atlas to create masks of 148 cortical regions (74 per hemisphere) (Destrieux et al. 2010) and Freesurfer’s automatic subcortical segmentation (ASEG) of 10 subcortical regions per hemisphere for 132 subjects. We calculated the average saliency value per region across all subjects (Fig. 3). Overall, the left hemisphere generally had a) higher magnitude of saliency value, with an average saliency of 6.07 × 10^-7^ ± 6.51× 10^-8^, while the average saliency of the right hemisphere was 1.96× 10^-8 7^ ± 2.76× 10^-9^ (region matched paired t-test: *p* = 1.00× 10^-14^, df = 73), and b) broader saliency range that was estimated by the average saliency range of each region in the left (1.99× 10^-7^ ± 2.95× 10^-8^) and right (8.60× 10^-9^ ± 1.76× 10^-9^) hemispheres across high and low severity subjects (region matched paired t-test: *p* = 9.14× 10^-9^, df = 73). In the left hemisphere, the highest magnitude of saliency appeared in the anterior and superior segments of the circular sulcus of the insula, the short and long insular gyri, the horizontal and vertical rami of the anterior lateral fissure, the orbital H-shaped sulcus, the orbital part of the inferior frontal gyrus, the inferior frontal sulcus, and the inferior precentral sulcus (Fig. 3). High magnitude saliency regions in the right hemisphere included the orbital H-shaped sulcus, anterior cingulate gyrus and sulcus, gyrus rectus, superior frontal gyrus, and subcallosal gyrus. To verify whether specific regions carrying the highest magnitude saliency were consistent across sites, we computed pairwise Jaccard index of the 30 highest magnitude regions (N=168 total regions) identified independently in each site-held-out fold. Mean pairwise Jaccard index was 0.37 (range 0.25-0.58) across folds (sites due to leave-one-site-out cross-validation), compared to the chance expectation of 0.10 for randomly selected sets of 30 regions. This, on average, corresponds to 16 out of 30 regions (Jaccard index of 0.37) shared between any two folds. Although regional saliency was not perfectly identical throughout folds, saliency regions with high magnitudes were more consistent across site-held-out folds than expected by chance.

To assess whether regional saliency reflected anatomical difference and not artifacts, we correlated each region’s average saliency with FreeSurfer-derived cortical thickness, gray matter volume, and surface area across 132 subjects and 148 regions (Table S5). Given the continuous and interval nature of these measures, Pearson correlation was used for this analysis instead of Kendall Tau. After Benjamini-Hochberg FDR correction, regional saliency was correlated with cortical thickness in 10 of 148 regions tested (r = 0.28-0.37), concentrated in the bilateral precentral, postcentral, and superior/middle frontal gyri and sulci. Saliency was correlated with gray matter volume in the postcentral gyrus (r = 0.30) and was not correlated with surface area in any region. Saliency of subcortical regions (20 regions) was not correlated with gray matter volume and mean intensity. This shows that saliency in sensorimotor regions and frontal cortex is related to structural variation in the cortex instead of arbitrarily distributed across the brain with no corresponding relationship in the subcortical regions.

Group-averaged saliency maps (Fig. 2) showed saliency from Integrated Gradients concentrated in the left lateral frontal cortex, with additional involvement in the anterior cingulate. Frontal and superior temporal regions carried high magnitude of saliency in both high- and low-severity groups. In low-severity subjects, the dorsolateral prefrontal and orbitofrontal cortex were both involved, whereas high-severity subjects showed saliency concentrated in the dorsolateral prefrontal cortex alone. Saliency was minimal in the right hemisphere across all groups (Fig. 2, Fig. 3). Standard deviation across subjects was substantial within the same left lateral frontal region that carried the highest magnitude of average saliency, reflecting between-subject variability in saliency magnitude.

**Figure 2:**
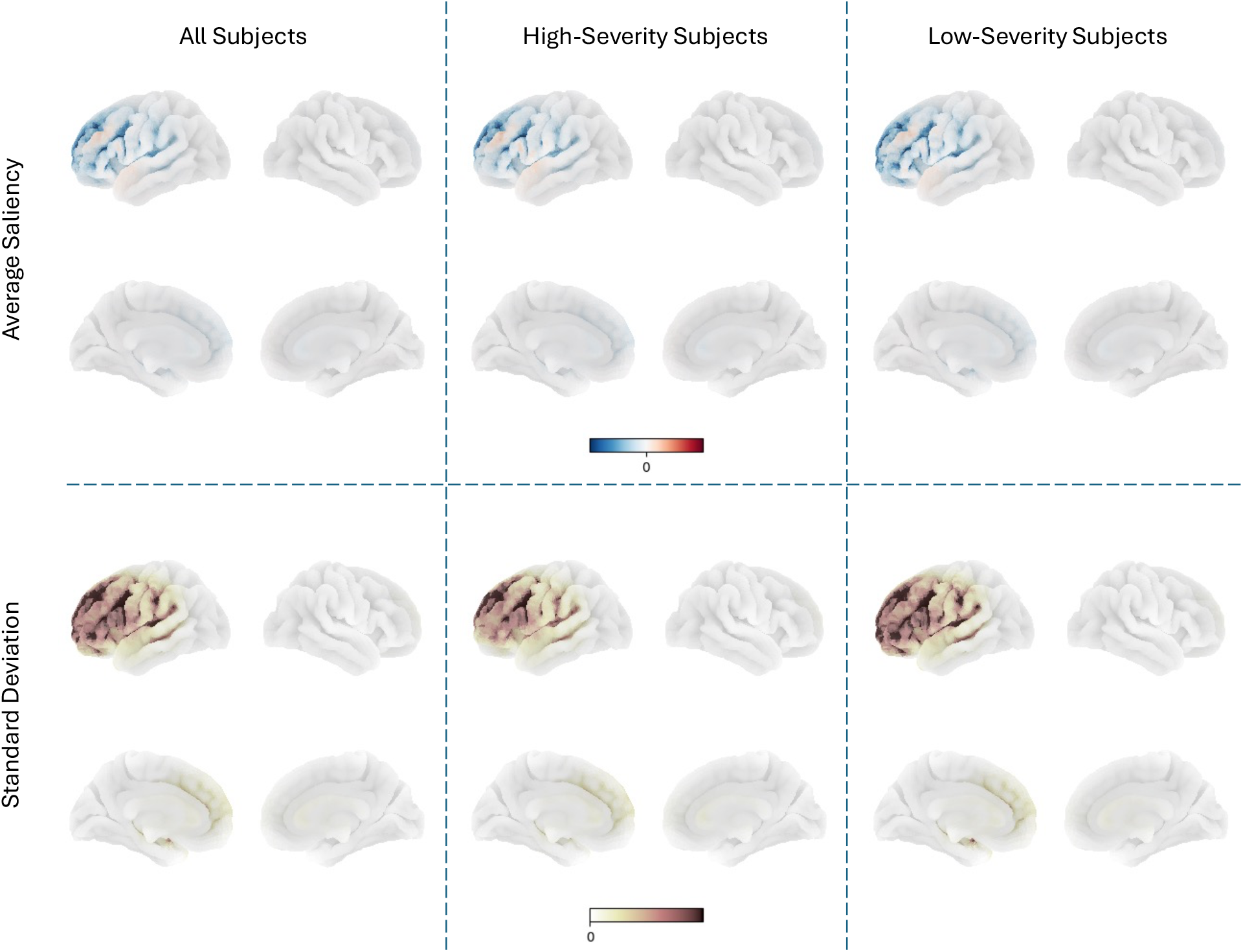
Group-averaged regional saliency and between-subject variability. Integrated Gradients saliency values were averaged across subjects and overlaid on FreeSurfer’s *fsaverage* pial surface in lateral (top) and medial (bottom) views for the left (LH) and right (RH) hemispheres. Columns show All Subjects (n=132), High-Severity Subjects (n=60), and Low-Severity Subjects (n=72). Top panels (Average Saliency): mean signed regional saliency per group (blue = negative, red = positive contribution to the model’s severity prediction). Bottom panels (Standard Deviation): standard deviation of regional saliency within each group on a sequential scale. Saliency was concentrated in left frontal, insular, and orbitofrontal regions across all three groupings, with minimal signal in the right hemisphere (see also Fig. 3). Between-subject variability was highest near sulcal regions, likely due to known large inter-subject sulcal differences in the frontal and temporal lobes (Vannier et al. 1991; Sanides 1962), and/or cortical folding abnormalities that have been reported in ASD (Nordahl et al. 2007).

**Figure 3:**
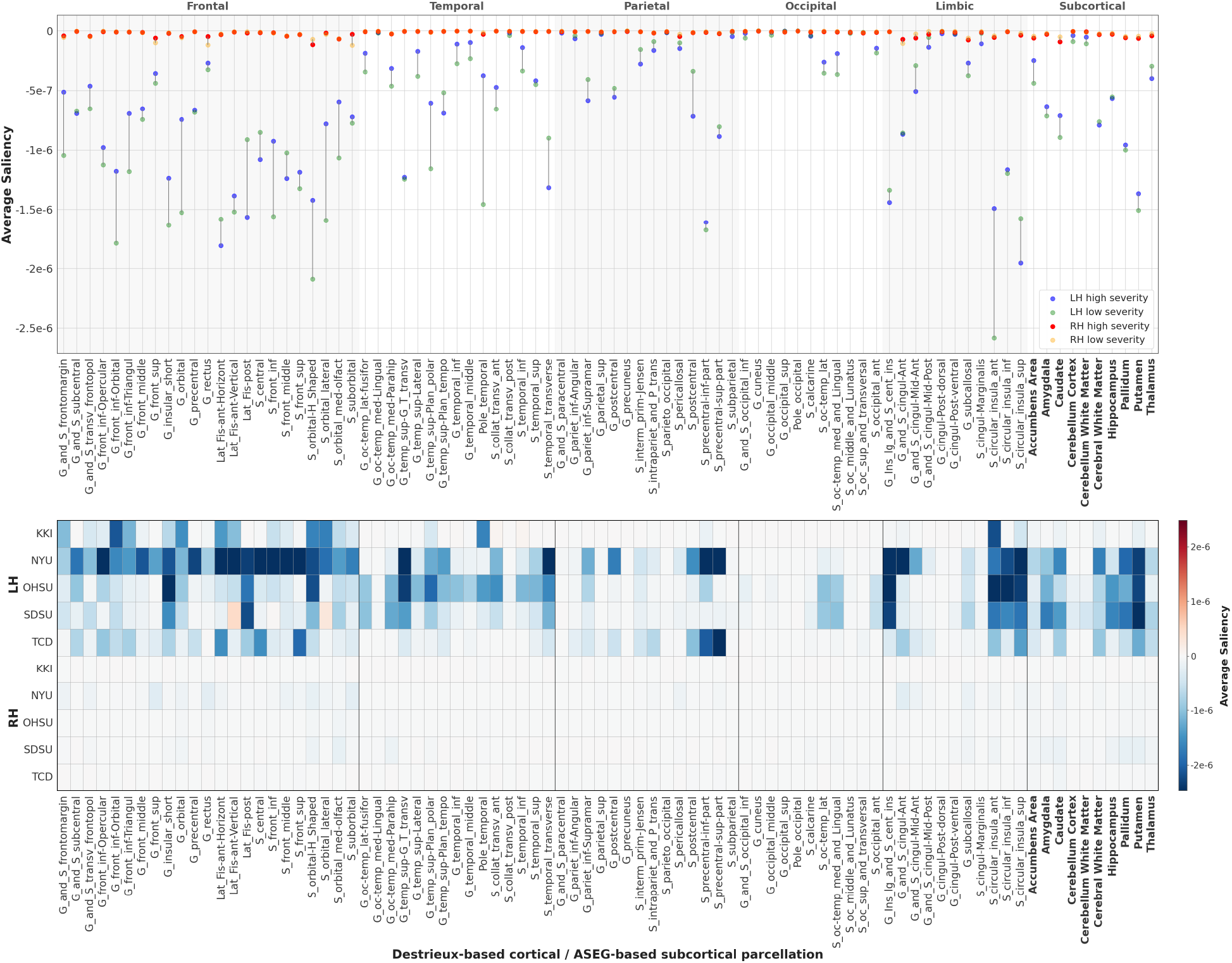
Regional saliency across cortical and subcortical parcellations. **Top:** average regional saliency between high- (blue) and low-severity (green) subjects in the left hemisphere (LH) and high- (red) and low-severity (yellow) in the right hemisphere (RH) for each cortical region labeled based on parcellation by Destrieux atlas and by FreeSurfer’s automatic subcortical segmentation of a brain volume (aseg atlas, bolded regions). Regions are grouped by lobe (Frontal, Temporal, Parietal, Occipital, Limbic) and subcortical structures. Note an overall larger magnitude and broader range of saliency in the left hemisphere. Bottom: average regional saliency per sMRI acquisition site (KKI, NYU, OHSU, SDSU, TCD) for LH (top 5 rows) and RH (bottom five rows).

### Covariation between regional saliency of cortical regions

Covariation between cortical regions can reflect coordinated importance that may be present in ASD. Kendall τ rank-based correlation between cortical regions in the Destrieux atlas were conducted to observe the relationship in paired saliencies within the brain. The top 1000 significant correlations sorted by the τ value are shown in Fig. 4. Each connecting thread indicates the presence of Benjamini-Hochberg FDR-corrected (*p_FDR* < 0.05) correlations with τ > 0.5. To maximize and optimize inference from these complex structural covariation maps, based on regional saliency correlations, we took the following steps. First, we used a stepwise adaptive threshold approach (from 0.1 to 0.7 with steps of 0.1) for examination of cross-regional correlations τ. We found that there were only few cross-regional correlations beyond threshold 0.8. The reason for the extensive, stepwise examination of τ thresholds, is because there were cortical regions that were significantly correlated with *many* other regions but with low τ, and regions correlated with a *few* other regions but with a higher τ. To distinguish between the two groups, we refer to the first as the regions *correlated with many* and the second, as the *highly correlated* regions groups, respectively. Both groups reflect extreme and in between scenarios that offer insights about potential regions that may act as potential network hubs in ASD. We generated Spring Cluster layout graphs (Fruchterman and Reingold 1991; Hamaguchi, Marumo, and Takeda 2025; NetworkX Developers 2025), which arranged cortical regions based on the number of significant cross-regional Kendall Tau correlations that each region maintained with other regions (Fig. 5). The algorithm created an interconnected network where each cortical region was a node, and significant correlations above the threshold became binary connections between regions. Regions were positioned such that those with a greater number of significant correlations with other regions were pulled toward the network center due to having more attraction forces acting upon them. This created a natural hierarchy, where regions serving as potential network hubs appeared in the center while regions with a smaller number of significant correlations appeared at the periphery. This spatial organization allowed identification of cortical regions that may function as potential structural pathology hubs in ASD, based on their tendency to show significant correlations with many other brain regions rather than the magnitude of individual correlation values. Fig. 5 shows the Spring Cluster layout with τ threshold of 0.6. At this threshold, the following regions were placed more toward the center, because they had a larger number of significant correlations with other regions: the left and right posterior rami of the lateral sulcus, the left lingual gyrus, the right superior segment of the circular sulcus of the insula, the left calcarine sulcus, the left intraparietal sulcus and transverse parietal sulci, the left superior parietal gyrus, the right long insular gyrus and central sulcus of the insula, the right short insular gyri, and the left supramarginal gyrus. For the full range of thresholds (0.1-0.7) to cover both groups of “correlated with many” and “highly correlated regions”, see Supplementary Table S7.

**Figure 4:**
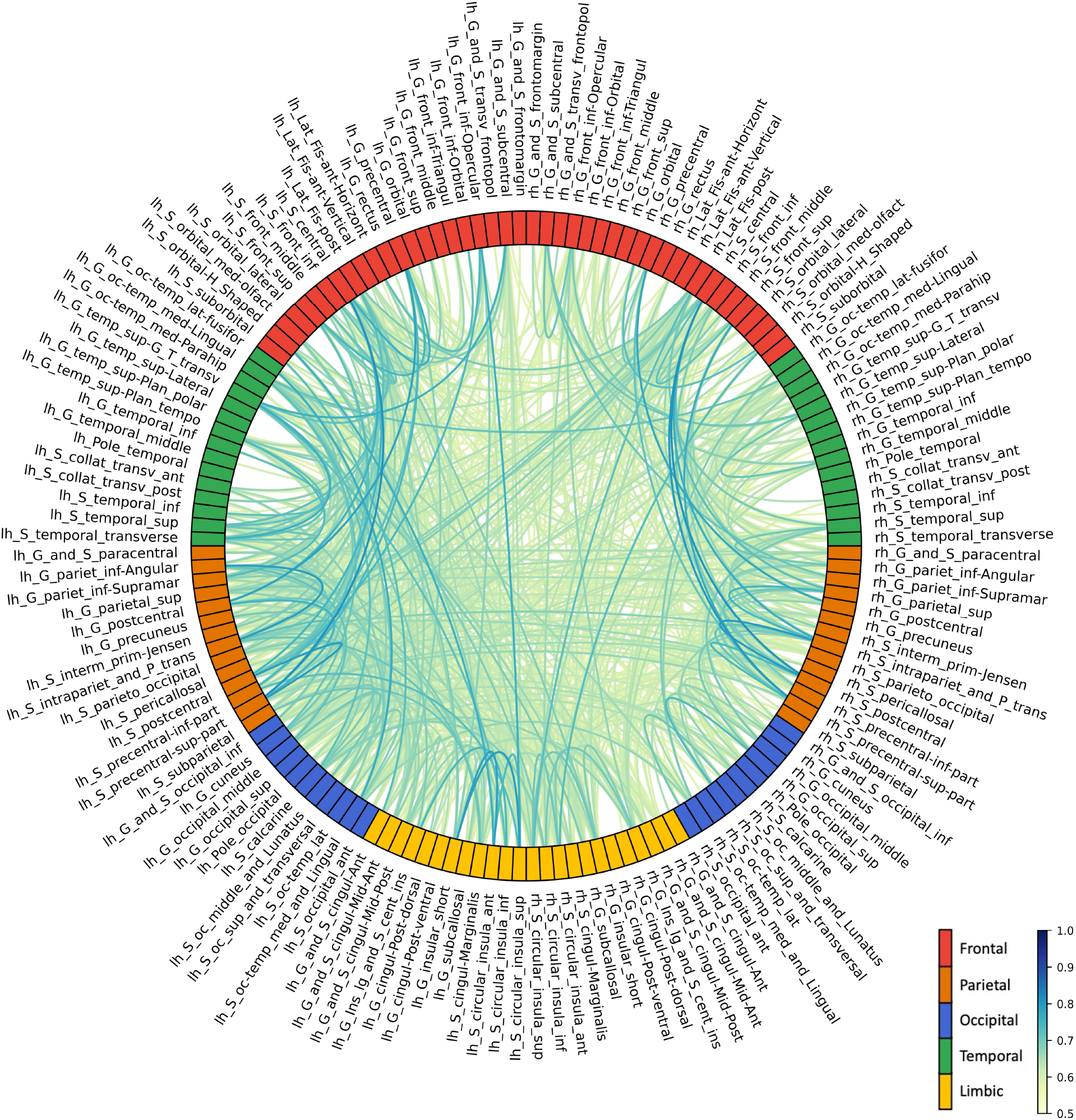
Circular covariation plot highlighting significant pairwise saliency correlations (Benjamini-Hochberg FDR-corrected *p* < 0.05 and where τ > 0.5) between cortical brain regions across high and low severity MRI cases. Darker lines represent higher Kendall τ whereas lighter lines represent lower Kendall τ values. The color of each brain region corresponds to the lobe the region is located in. The strongest correlations were mostly between regions within the same lobe.

**Figure 5:**
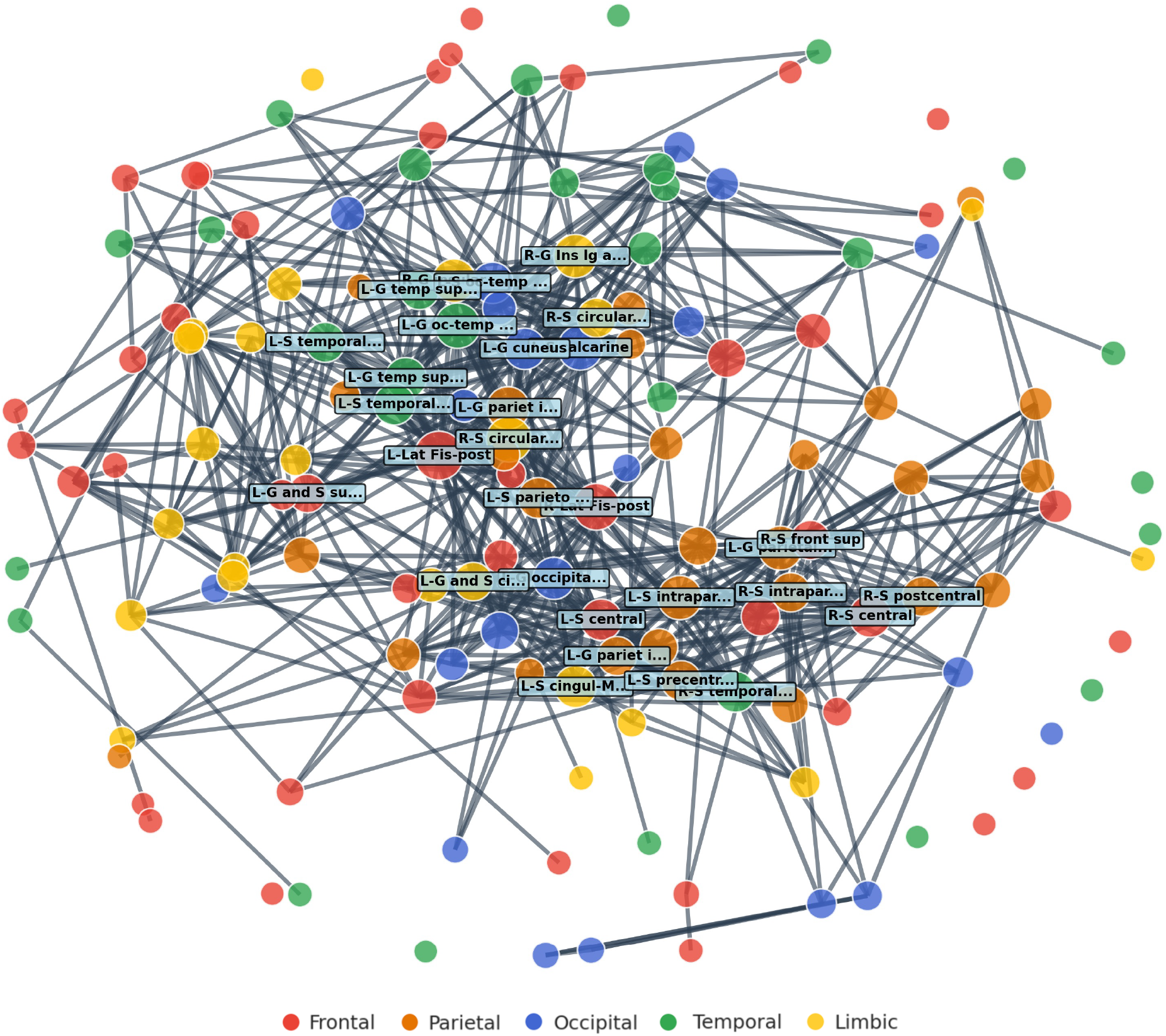
Spring Cluster layout graph for corticocortical saliency correlations. Covariation brain regional analysis dependent on region-region Tau correlations depicted in Figure 4. Each node represents a region in the Destruiex atlas, highlighted with their corresponding lobe color. Size of each node is dependent on how many significant correlations (Benjamini-Hochberg FDR-corrected *p* < 0.05 and where τ > 0.6) the respective region takes part in. The distance between correlated regions is weighted by 1– τ, where highly correlated regions are closer together. The algorithm arranges the nodes to minimize the weights between the regions. This leads to clustering of highly correlated regions (top 30 regions are named) towards the center and less correlated regions to the periphery.

To determine whether the regional saliency covariation network (Figs. 4-6) reflected anatomical organization, we performed an inter-structural-covariance-analysis (ISCA) to compare the saliency covariation network to structural covariance networks built from cortical thickness, gray matter volume, and surface area (Mantel test, 2000 permutations). Saliency covariation was found to be associated with all three structural covariance networks across all subjects and within high- and low-severity groups after Benjamini-Hochberg FDR correction (Mantel r = 0.11-0.26, *p* ≤ 0.014) (Table S6). This shows that saliency covariation reflects covariance found across morphologic metrics.

To better understand the organization and likely functional effects of relevant cortical regions that showed a high number of correlations with other regions or high correlation values in ASD, we counted the number of relevant cortical regions belonging to 4 main functional brain networks, consistently affected in ASD, i.e., Default Mode (DMN), Salience (SN), Frontoparietal (FPN, or central executive network), and Social networks, as well as 4 additional functional brain networks, i.e., Language, Sensorimotor, Dorsal/Ventral attention, and Limbic/Emotion networks, which have been associated with ASD dysfunction. For example, using τ = 0.5, Fig 6A pie chart shows the relative involvement of the 8 functional brain networks, highlighting the relative representation of regions that are part of each of these 8 functional brain networks, which are well-established in ASD pathology. The raw counts, such as 26 for the Dorsal/Ventral Attention Network is the count of brain regions belonging to Dorsal/Ventral Attention Network in Table S7 with threshold τ = 0.5, and the ratio 26/106 = 24.5% indicates the ratio of DMN involvement in which 106 is the total brain region count across all 8 functional brain networks in the pie chart.

**Figure 6:**
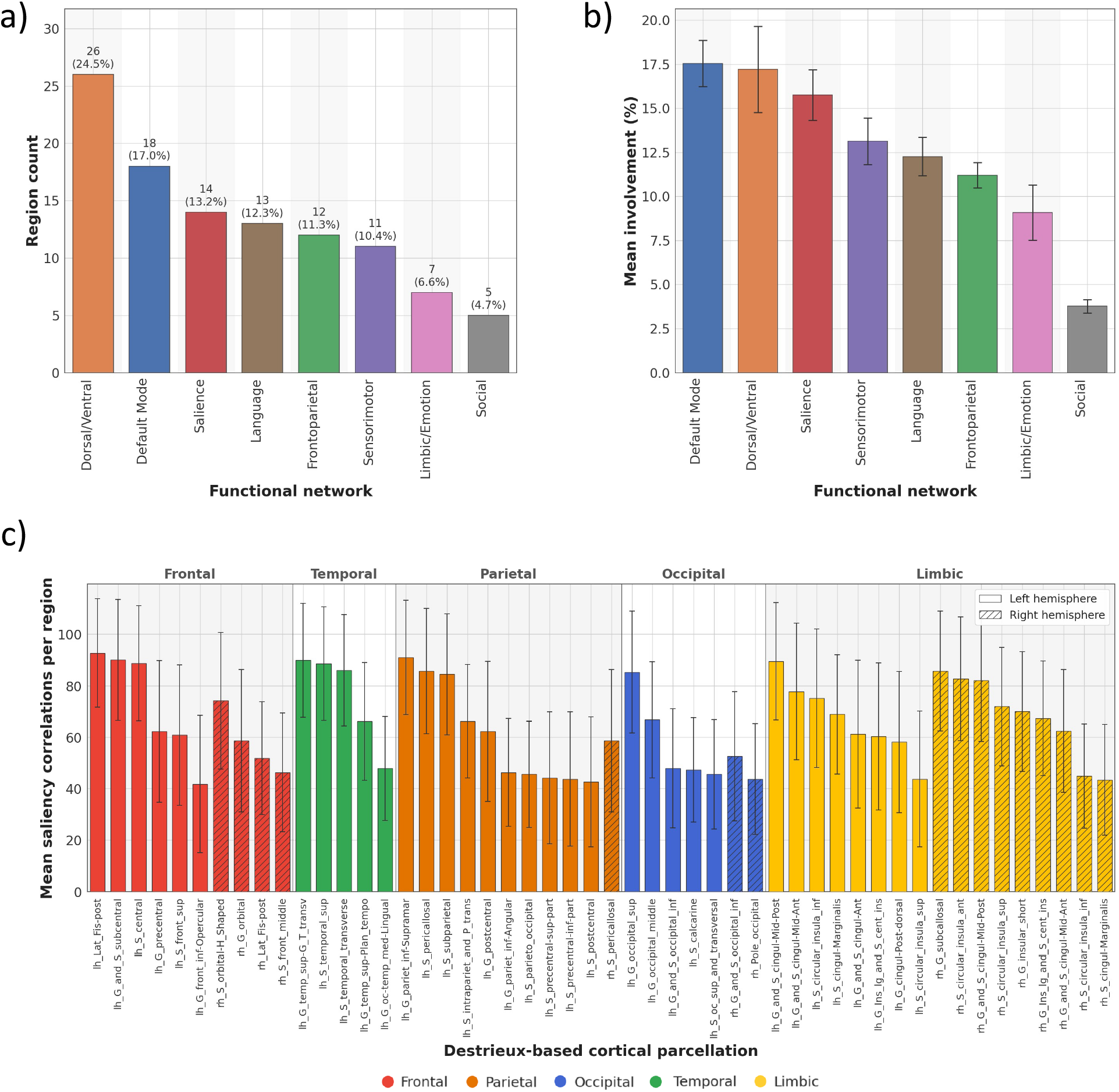
Degrees of involvement of functional brain networks and lobes in ASD. **A**. The degree of involvement for each brain network was derived from the pairwise brain region saliency correlation analysis shown in Fig. 4. Specifically, we considered significant correlations (Benjamini-Hochberg FDR-corrected *p* < 0.05) with τ exceeding a set threshold (e.g., 0.5) and selected the top *n* brain regions (e.g., 50) with the highest number of such correlations. From this list, we counted how many regions belonged to each functional network (raw counts shown beside the bar chart) and calculated the proportion of each network’s count relative to the total, expressed as a percentage. **B**. The analysis in **A** was repeated and averaged across thresholds ranging from 0.1 to 0.7 in increments of 0.1 (error bars represent sem). These measures indicate the global engagement of regions within functional networks with the rest of the brain in ASD. **C**. The number of correlations per brain region was summed across the top 50 areas from each threshold-based table in S3, then divided by seven (the number of τ thresholds between 0.1 and 0.7) to calculate the average number of correlations per region across all thresholds (error bars represent sem). Areas were subsequently re-ranked based on these averaged values to identify the top 50 regions involved in ASD according to averaged structural correlations. The largest number of regions involved in ASD are found in the frontal and temporal lobes in the left hemisphere.

For the overall examination of functional brain network involvement across the full range of τ thresholds (0.1-0.7), we averaged the brain region counts and their ratios per network across the 7 pie charts, similar to the one in Fig 6A, for the 7 thresholds, 0.1-0.7 (Fig. 6B, Table S7). Overall, the Dorsal/Ventral attention network and Default Mode Networks consistently showed the highest average representation and association with ASD, among functional networks, followed by the Salience, Sensorimotor, Language, Frontoparietal, and Limbic/Emotion networks. The Social networks were minimally represented, reflecting their more distributed or overlapping nature in the anatomical brain regions.

Additionally, we summed the number of correlations per brain region across the top 50 areas from each threshold-based table in Supplementary Table S7 and divided by 7 (number of τ thresholds between 0.1-0.7) to reflect the average number of correlations per brain region across tables covering all thresholds. We then, re-ranked the areas based on these averaged correlations to identify the top 50 regions involved in ASD based on averaged structural correlations (Fig. 6C). This analysis showed that pairwise saliency correlation across cases was dominant in the left hemisphere, both in terms of number of inferred structural covariation per region (higher values in the average covariation columns) and in terms of the number of regions as potential central hubs (more table rows for left than right).

### Correlations regarding regional saliency averages

We investigated correlations between average and standard deviation of saliency within either LH or RH regions from cases either with high or low severity, resulting in an 8×8 grid accounting for LH vs RH (2) × Average (Avg) vs Standard deviation (STD) (2) × High vs Low severity (2) scatter plots and distribution histograms (Fig. 7). Overall, the patterns of distribution and scatter indicate more proportional area saliency average and standard deviation in LH in high severity cases, less proportionality in RH in high severity, and no proportionality of area saliency average in one hemisphere and standard deviation of equivalent area in opposite hemisphere in high and low severity cases. This comparison showed that LH structural modifications in high severity cases generate STD proportional to Avg saliency across subjects, indicative of proportional modifications across regions, such as inflated or compressed saliency. The degree of proportionality was smaller in low severity LH and high and low severity RH, and nonexistent across LH and RH. Together, these findings revealed systematic heterogeneity, especially pronounced in LH in high severity ASD and indicate that the heterogeneous structural impact of ASD on each left-brain region was proportional to the mean structural impact on that left brain region across affected individuals. In this regard, the structural saliency of all regions within each lobe in the LH for high and low severity were not normally distributed (see diagonal panels with distribution plots in Fig. 7), especially compared to the right hemisphere. This strongly suggests that structural aberrations in LH regions were not an artifact of random distribution across MRI scans and there were macro-structural effects reflected in MRI saliency. This is consistent with the dominant involvement of left hemisphere in ASD.

**Figure 7.**
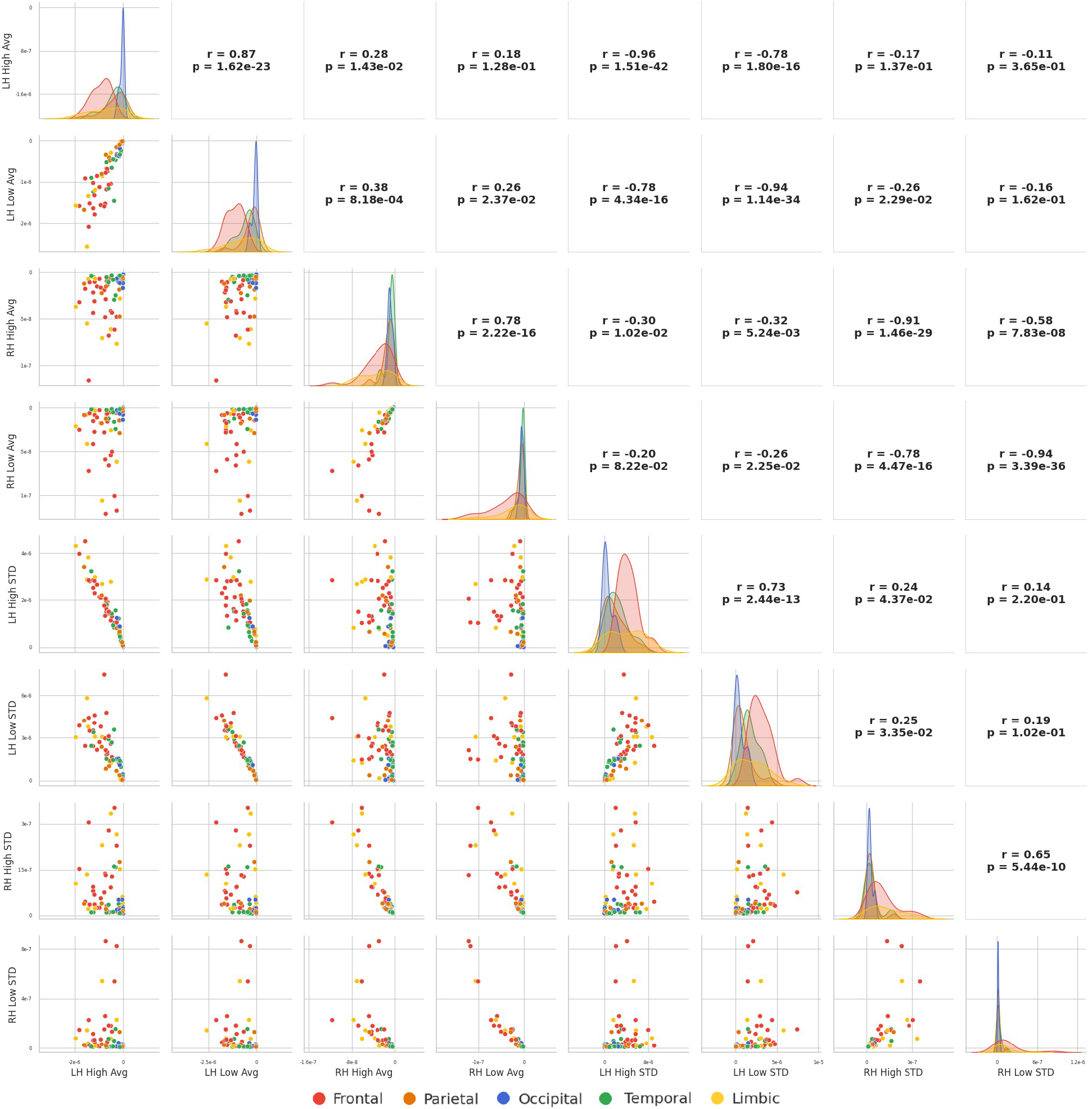
Scatter plots and distributions of the average regional saliency and the associated standard deviation for each cortical brain region. Plots are organized on an 8×8 grid, based on specifications in Fig 3. Each point in the scatter plot belongs to a brain region from Destrieux atlas similar to Fig. 3, total of 74 regions. Scatter plot axes are restricted to the left (LH) or right (RH) hemisphere, high (High) or low (Low) severity score cases, and the metrics include average (Avg) and standard deviation (STD) across the targeted cases (high or low severity). For example, an axis label of LH High STD refers to regions in the left hemisphere of all high severity cases averaged for each brain region. Therefore, each scatter plot has 74 points corresponding to one of the 74 brain regions listed in Fig. 3. Brain regions belonging to each lobe are color coded in all scatter and distribution (diagonal of the grid) plots. Key findings include a) maximum trend of *linear* scatter within LH high score, less in LH low score, RH high and low scores, and isotropic or no trend scatter across LH and RH (either with high or low score), b) distribution plots on the grid diagonal are more normal like for RH and low severity and max deviated from normal for LH high severity. For interpretations, see the text.

### Correlating regional saliency and ASD severity across cases

We analyzed correlations using Kendall rank correlation coefficient τ between the SRS subscores (cognition, awareness, communication, mannerisms, and motivation) and the saliency of each brain region across low to high severity ASD cases. For these analyses, we considered 148 regions (2×74 for both left and right hemispheres) labeled using the Destrieux atlas parcellation and found that cognition, awareness, communication, mannerisms, and motivation subscores were nominally (*p* < 0.05) correlated with a distinct number of cortical regions in each lobe of the left and right hemisphere (3/1, 1/0, 5/10, 2/4, 5/2, respectively; Table 1), however, did not survive FDR correction (Table S8). Correlations in the left hemisphere with SRS subscores primarily featured regions from the temporal (6 correlations with SRS subscores involving 3 regions) and parietal (8 correlations with SRS subscores involving 4 regions) lobes while correlations in the right hemisphere featured regions in the occipital lobe (10 correlations with SRS subscores involving 5 regions).

**Table 1.** Brain regions associated with each ASD sub-score, grouped by hemisphere and lobe.

| Sub-score | Left Hemisphere | Right Hemisphere |
| --- | --- | --- |
| Cognition | **Frontal:** Orbital gyri.<br>**Temporal:** Planum polare of the superior temporal gyrus.<br>**Parietal:** Primary intermediate sulcus. | **Occipital:** Medial occipito-temporal and lingual sulci. |
| Awareness | **Parietal:** Primary intermediate sulcus. | None |
| Communication | **Temporal:** Temporal plane of the superior temporal gyrus.<br>**Parietal:** Supramarginal gyrus, Primary intermediate sulcus, Pericallosal sulcus.<br>**Occipital:** Lateral occipito-temporal sulcus. | **Temporal:** Inferior temporal gyrus, Anterior transverse collateral sulcus.<br>**Parietal:** Precuneus gyri, Pericallosal sulcus, Subparietal sulcus.<br>**Occipital:** Cuneus, Calcarine sulcus, Lateral occipito-temporal sulcus, Medial occipito-temporal and lingual sulci. |
| Mannerism | **Temporal:** Posterior transverse collateral sulcus.<br>**Parietal:** Primary intermediate sulcus. | **Temporal:** Anterior transverse collateral sulcus.<br>**Parietal:** Subparietal sulcus.<br>**Occipital:** Calcarine sulcus, Lingual part of the medial occipito-temporal gyrus. |
| Motivation | **Temporal:** Planum polare of the superior temporal gyrus, Temporal plane of the superior temporal gyrus, Temporal Pole.<br>**Parietal:** Supramarginal gyrus, Primary intermediate sulcus. | **Temporal:** Anterior transverse collateral sulcus.<br>**Occipital:** Medial occipito-temporal sulcus and lingual sulcus. |

## Discussion

Advances in genetic, molecular, histopathological, and imaging studies, as well as clinical diagnosis of neurodevelopmental, neurodegenerative, and mental disorders, provide a wealth of data that can facilitate fine quantitative characterization of their heterogenous nature. Combined with major developments in the field of explainable AI, through techniques that can highlight the salient input features contributing to classification, make it possible to systematically interrogate and identify the underlying structural and functional correlates of such heterogeneities Within this framework, in this study we developed explainable AI approaches and applied these to study *in vivo* structural MRI sequences from individuals with ASD from the ABIDE II dataset with the goal to correlate key structural changes in the autistic brain with the functional, cognitive, and behavioral heterogeneity of ASD. We parsed heterogeneity through a) correlations between the phenotype spectrum, in terms of diagnostic subscores, with the AI derived spectrum of structural and anatomical saliency, and b) pairwise correlations between the AI derived spectrum of structural saliency as a pointer to co-involvement and functional brain network changes in ASD. Using this approach, we trained a pretrained MedicalNet backbone with frozen weights and a trainable classification head to distinguish between high- and low-severity ASD subjects from sMRI. This approach limited the number of parameters to prevent overfitting and was validated by leave-one-site-out cross-validation, label-permutation testing, and seed stability. Based on the validated 3D CNN, we extracted relevant brain regions that contributed to the classification and measured a) the correlation of their saliency level with the cognitive and behavioral heterogeneity, and b) their pairwise saliency correlation as pointers to brain functional networks and their heterogenous modifications in ASD. We then extracted the regions of interest, mapped through cortical and subcortical segmentation, by generating MRI saliency maps using a gradient-based method.

Our findings about brain structural changes in ASD can be summarized as a) differential involvement of left vs right hemispheres with more involvement of left hemispheric regions and functional networks, b) more extent (correlation count) in the ASD involvement of the left brain, c) more left brain structural changes correlated with ASD subscores and phenotype, and therefore d) more reflection of ASD heterogeneity in left brain correlated with phenotypical and structural changes. Among brain lobes and functional networks, the frontal and temporal lobes and the default mode, central executive, and dorsal/ventral attention networks showed most structural changes in ASD. Our systematic Al approach leveraged the phenotypical heterogeneity and spectral nature of ASD to uncover structural brain region heterogeneity in ASD, as well as correlations between areas and across structural and phenotypical/diagnostic heterogeneity. This revealed global, regional, and network changes in ASD associated with symptom severity and diagnostic scores.

The constellation and configuration of high vs. low saliency and correlations we found, revealed many functional networks and areas that are affected in ASD, including but not limited to multiple brain regions that are posterior components of the default mode network, bilateral precuneus, right inferior occipital gyrus, left cerebellar regions (Liloia et al. 2023). Significant positive or negative structural saliency correlations revealed in this study can be in part the origin or outcome of functional connectivity changes (over- or under-connectivity) in ASD, together motivating the consideration of brain structural phenotypes of ASD and laying a framework for neuroanatomical heterogeneity in ASD (Shan et al. 2022). Overall, ASD exhibited heterogenous and widespread structural and functional feature prints. The saliency overlay results, showed distribution across areas and hemispheres with high saliency in frontal, limbic and temporal areas. Areas with a role in speech, language, comprehension, and non-verbal communication displayed high saliency. These included pars opercularis, pars orbitalis, inferior frontal regions (including Broca’s area), superior temporal sulcus (Bigler et al. 2007; Foundas et al. 1996; Peeva et al. 2013; Yamasaki et al. 2010), and the horizontal, vertical, and posterior rami of the lateral sulcus, which define the pars triangularis, pars orbitalis, and pars opercularis regions and also hold and Wernicke’s area (Chaichana and Quiñones-Hinojosa 2019; Griffiths et al. 2010). Other areas with high saliency included the insula that has a role in sensory integration and regulation of emotion (Viñas-Guasch and Wu 2017), and the temporal pole, which is commonly associated with visual processing, socio-emotional processing, and other cognitive processes (Herlin, Navarro, and Dupont 2021). Interestingly, left hemisphere regions throughout most of the cortex showed higher saliency values and larger separation across the spectrum of low-high ASD severity whereas regional saliency in the right hemisphere remained lower in magnitude and with smaller separation across the ASD spectrum.

The saliency-based correlations highlighted involved functional networks, covariation and facilitated ranking of regions in terms of affected correlations. In line with prior reports [reviewed in (Amaral et al. 2008)], our findings also showed the heterogeneous and distributed neuroanatomy of ASD (Amaral et al. 2008), which is associated with functional connectivity and network aberrations (Müller 2007), including widespread reported over- and under-connectivity (Baran et al. 2023; Cherkassky et al. 2006; Just et al. 2007; Kenet et al. 2012; Khan et al. 2013; Müller 2007). Our analysis revealed functional networks implicated in ASD, with the default mode (DMN), central executive (CEN or frontoparietal), dorsal/ventral attention, sensorimotor, and salience (SN), networks at the top of the list. These findings are in line with previous studies by Uddin et al. (2013) that also highlighted the involvement of these and other functional networks, co-involvement, and regions in ASD. Further support for our findings comes from observations of weaker resting state functional connectivity across distributed functional networks in ASD, including frontoamygdala functional connectivity that likely contributes to socioemotional impairments (Odriozola et al. 2019). Resting state underconnectivity has been reported in functional studies using the ABIDE dataset (Duan et al. 2017), indicating underconnectivity in areas within the default mode network (DMN) and within the ventral attention network (VAN), as well as between pairs of networks, including DMN and Visual Network (VN), dorsal attention network (DAN) and VAN, and limbic network (LN) and frontoparietal network (FPN). Schipul et al. (Schipul et al. 2012) showed decreased functional and structural connectivity measures consistent with the corpus collosum size change in ASD compared to TD, which could account for some of the hemispheric differences we observed. Functional and anatomical underconnectivity has also been noted by (Just et al. 2007), using functional measures (synchronization) between parietal and frontal areas in proportion with the smaller size of the genu of corpus callosum. Local and long-range functional connectivity was shown to be reduced within FFA, and between the FFA and two distant cortical regions, the left inferior frontal gyrus (IFG) and left anterior cingulate cortex (Khan et al. 2013), which were also relevant in our analysis. The anterior cingulate cortex, in particular, has been a consistent site of ASD pathology in imaging (Thakkar et al. 2008) and histopathological and computational studies (Yazdanbakhsh et al. 2025; Zikopoulos, Liu, et al. 2018; Zikopoulos and Barbas 2010; Zikopoulos, García-Cabezas, and Barbas 2018). Moreover, several resting-state functional networks may be more loosely connected in ASD, based on lower covariation of the anterior cingulate region with posterior cingulate and precuneus areas, between anterior and posterior cingulate, and lower functional connectivity between anterior and posterior medial cortices (Cherkassky et al. 2006). In addition, Ecker et al. 2010 identified distributed functional networks, limbic, frontal-striatal, fronto-temporal, fronto-parietal and cerebellar systems that distinguished ASD from controls, with better gray matter-compared to white matter-based classification. On the other hand, regions and functional networks that were relevant in our analyses have also been implicated in reports of hyperconnectivity in the salience network (SN), composed of anterior cingulate cortex ACC, ventral anterior insular cortex, amygdala, hypothalamus, ventral striatum, thalamus, and brainstem nuclei (Uddin et al. 2013), as well as social network-related long- and short-range hyperconnectivity (Supekar et al. 2013).

## Methodological considerations and limitations

In this study we used the 3D ResNet-50 allowed for the whole MRI volume to be used for training. Although 3D CNNs can analyze data across MRI slices rather than the data on the slice itself, they are extremely prone to variability in data. It was therefore essential, for accurate and consistent performance of the 3D neural network, to reduce variability through MRI preprocessing.

Importantly, our study focused on male subjects, due to limited availability of MRI sequences and accompanying diagnostic/phenotypic data for females therefore, reflecting heterogeneity also seen in other studies of male individuals (Chen et al. 2019). Moreover, the amount of publicly available MRI datasets that include phenotypic data such as the SRS Total and its subscores is limited, which naturally could make our model prone to overfitting, impacting extracted regions of interest, based on saliency maps and saliency magnitude. Hence, why we used leave-one-site-out, seed stability, label-permutation test, model randomization, saliency seed stability, and ISCA to make sure that saliency was representable of ASD.

IQ differed across high- and low-severity groups after FDR correction (p_FDR = 8.1× 10^-5^), which is consistent with prior work showing an inverse association between ASD symptom severity and IQ across large samples (Mazurek et al. 2019; Denisova and Lin 2023). As we did not directly test whether IQ is encoded in or contributes to CNN’s classification or saliency patterns, we cannot completely distinguish severity-specific structural signal from IQ related structural variation in this analysis.

Additionally, the saliency maps depict the contribution of each voxel to making the low to high severity ASD prediction, as a macrostructural pointer to ASD, and the highly significant saliency correlations across pairs of regions can only suggest their involvement in ASD as part of brain functional networks. Nuances such as intra-areal and inter-areal hypo- and hyper-connectivity changes in ASD (Cherkassky et al. 2006; Just et al. 2007; Kenet et al. 2012; Khan et al. 2013; Müller 2007; Supekar et al. 2013) appear as a tip of iceberg in saliency correlation analysis, and neuronal, connectional and neurobiological changes in ASD demands approaches with much higher spatial resolution. Therefore, the current study can at best outline underlying brain region and functional network heterogeneity in the severity of ASD.

## Conclusions

Our findings showed that structural MRI contains information that reflects functional network and diagnostic score changes in ASD, which can be identified, using state-of-the-art AI, to pinpoint relevant brain regions and their functional networks underlying ASD heterogeneity. Our analyses provided distinct involvement measures of frontal, temporal, parietal, and occipital lobes, and limbic, insular, and subcortical regions and their constituent areas and subregions in social-awareness, attention, social interactions, and cognitive control scores. Additional correlation and subsequent ranking revealed hierarchically arranged involvement of functional networks in ASD, which, in descending order of impact, included default mode (DMN), dorsal/ventral attention, central executive (CEN or frontoparietal), sensorimotor, salience (SN), language, limbic/emotion networks, and social. These findings highlight novel, detailed, and systematic anatomical, functional, and phenotypical changes and associations that can be used as biomarkers to parse and frame the heterogeneity and spectral nature of ASD.

