## Supplemental material for "Structural changes in autism reflect atypical brain network organization and phenotypical heterogeneity: a deep network approach"

### Supplementary Materials

| ABIDE-II Site | Total Subjects | Low Severity Subjects | High Severity Subjects | Total Subjects Used |
|---|---|---|---|---|
| **NYU-1** | 72 | 15 | 22 | 37 |
| **SDSU** | 55 | 6 | 11 | 17 |
| **TCD** | 41 | 2 | 2 | 4 |
| **OHSU** | 79 | 11 | 7 | 18 |
| **KKI** | 147 | 41 | 20 | 61 |

**Table S1:** Number of subjects acquired from each ABIDE-II site and the split of low and high severity subjects used per site. Total subjects refer to the total number of subjects each site provides to the ABIDE-II database whereas, total subjects used refers to the number of subjects that fit the criteria of male, age 5 to 12 and fit the SRS Total threshold. Note that some subjects are not used in analysis to have a balanced low severity to high severity subject ratio.

| Site | Scanner | Sequence | TR (ms) | TE (ms) | TI (ms) | Flip Angle (°) | Acquired Voxel (mm) | Matrix | FOV (mm) | Slices / Thickness | Bandwidth |
|---|---|---|---|---|---|---|---|---|---|---|---|
| **NYU 1** | Siemens Allegra 3T | MPRAGE | 2530 | 3.25 | 1100 | 7° | 1.3 × 1.0 × 1.3 | Base res. 256, 75% phase res. | 256 (read), 100% phase | 128 / 1.33 mm | 200 Hz/px |
| **KKI (8-ch)** | Philips 3T | 3D TFE | 8 | 3.7 | Not reported | 8° | 1.00 × 1.00 × 1.00 | 256 × 200 | 256 × 200 × 200 | 200 / 1.0 mm | 191.5 Hz/px |
| **KKI (32-ch)** | Philips 3T | 3D TFE | 8.2 | 3.7 | Not reported | 8° | 1.00 × 1.00 × 1.00 | 212 × 172 | 172 × 212 × 150 | 150 / 1.0 mm | 192.9 Hz/px |
| **TCD** | Philips 3T | 3D TFE | 8.4 | 3.9 | Not reported | 8° | 0.90 × 0.90 × 1.80 | 256 × 256 | 230 × 230 × 162 | 180 / Not separately reported† | 188.3 Hz/px |
| **OHSU** | Siemens TrioTim 3T | MPRAGE | 2300 | 3.58 | 900 | 10° | 1.0 × 1.0 × 1.1 | Base res. 256, 100% phase res. | 256 (read), 93.8% phase | 160 / 1.10 mm | 180 Hz/px |
| **SDSU** | GE 3T | 3D SPGR (FSPGR) | Not reported | Not reported | 600 | 8° | Not reported§ | Freq 256 / Phase 192 | 256 (25.6 cm) | 176 / 1.0 mm | Not reported |

**Table S2:** Scanner, repetition time (TR), echo time (TE), inversion time (TI), flip angle, acquired voxel, matrix, field of view, slices / thickness, bandwidth, and scan time for ABIDE-II sites used in the study (NYU, KKI, TCD, OHSU, and SDSU).

| Variable | Low severity mean ± SD | High severity mean ± SD | Low severity n | High severity n | Test | Statistic | p | p_FDR |
|---|---|---|---|---|---|---|---|---|
| **eTIV** | 1,511,017.19 ± 123,146.42 | 1,496,198.00 ± 140,153.50 | 72 | 60 | t-test | -0.6389 | 0.524149 | 0.524149 |
| **IQ** | 116.10 ± 11.67 | 105.15 ± 15.32 | 72 | 59* | t-test | -4.5171 | 0.000016 | 0.000081 |
| **Age** | 9.67 ± 1.47 | 9.25 ± 1.69 | 72 | 60 | Mann-Whitney U | 1835.5 | 0.138639 | 0.173299 |
| **Euler total** | −201.94 ± 97.63 | −284.60 ± 140.47 | 72 | 60 | Mann-Whitney U | 1255 | 0.000036 | 0.000089 |
| **Site** | NA | NA | NA | NA | Chi-square (df=4) | 9.8169 | 0.043628 | 0.072713 |

**Table S3**: Distribution of confound variables (estimated total intracranial volume, IQ, age, Euler number, and site) by severity group (mean ± SD, n), test, test statistic, and p value (nominal and Benjamini-Hochberg FDR corrected).

| Fold (Site) | Mean Balanced Acc (1000 runs) | Low | High | Lowest Val Loss | Accuracy at Lowest Val Loss | Run # (lowest val) |
|---|---|---|---|---|---|---|
| KKI_1 | 0.5391 | 0.399 | 0.626 | 0.5463 | 0.5402 | 659 |
| NYU_1 | 0.5054 | 0.426 | 0.603 | 0.615 | 0.4712 | 791 |
| OHSU_1 | 0.6962 | 0.273 | 0.812 | 0.5275 | 0.7662 | 928 |
| SDSU_1 | 0.578 | 0.303 | 0.735 | 0.6228 | 0.6515 | 95 |
| TCD_1 | 0.7275 | 0.25 | 1 | 0.5986 | 0.75 | 490 |
| **Mean across folds** | **0.6092** | **0.3302** | **0.7552** | | **0.6358** | |

**Table S4**: Classification performance from leave-one-site-out cross-validation across 1000 seed stability runs. Mean balanced accuracy, lowest accuracy, highest accuracy, lowest validation loss, accuracy at lowest validation loss, and run number at lowest validation loss reported per site. Integrated Gradients were computed using the model weight using the model with the lowest validation loss per site.

| Region | Gray matter volume | Surface area | Cortical thickness |
|---|---|---|---|
| **lh-G&S_cingul-Ant** | -0.0536 | 0.0395 | -0.0174 |
| **lh-G&S_cingul-Mid-Ant** | -0.1410 | -0.0635 | -0.0997 |
| **lh-G&S_cingul-Mid-Post** | -0.0940 | -0.0541 | -0.1457 |
| **lh-G&S_frontomargin** | 0.2711 | 0.1044 | 0.1419 |
| **lh-G&S_occipital_inf** | 0.0271 | -0.0168 | 0.0780 |
| **lh-G&S_paracentral** | -0.0287 | -0.1292 | 0.1247 |
| **lh-G&S_subcentral** | -0.1200 | -0.0832 | 0.0182 |
| **lh-G&S_transv_frontopol** | 0.1256 | -0.0645 | 0.1874 |
| **lh-G_Ins_lg&S_cent_ins** | 0.0062 | 0.0226 | -0.0027 |
| **lh-G_cingul-Post-dorsal** | -0.1203 | -0.1235 | -0.0959 |
| **lh-G_cingul-Post-ventral** | 0.0980 | 0.1146 | 0.0179 |
| **lh-G_cuneus** | 0.2630 | 0.1261 | 0.2004 |
| **lh-G_front_inf-Opercular** | 0.0208 | 0.0238 | -0.0208 |
| **lh-G_front_inf-Orbital** | 0.0257 | 0.0993 | -0.0773 |
| **lh-G_front_inf-Triangul** | 0.0357 | 0.0267 | -0.0050 |
| **lh-G_front_middle** | 0.1503 | -0.0042 | **0.3471** |
| **lh-G_front_sup** | 0.1559 | 0.0215 | 0.2532 |
| **lh-G_insular_short** | -0.0541 | 0.0178 | -0.0656 |
| **lh-G_oc-temp_lat-fusifor** | -0.0569 | -0.1299 | 0.1400 |
| **lh-G_oc-temp_med-Lingual** | 0.0798 | 0.1092 | -0.0691 |
| **lh-G_oc-temp_med-Parahip** | 0.0465 | 0.0336 | 0.1628 |
| **lh-G_occipital_middle** | 0.0671 | 0.0326 | 0.0854 |
| **lh-G_occipital_sup** | -0.0411 | -0.0109 | 0.0394 |
| **lh-G_orbital** | -0.0051 | 0.0737 | -0.0995 |
| **lh-G_pariet_inf-Angular** | -0.1646 | -0.0926 | -0.0467 |
| **lh-G_pariet_inf-Supramar** | -0.0768 | -0.0795 | 0.1236 |
| **lh-G_parietal_sup** | 0.0784 | 0.0822 | 0.0376 |
| **lh-G_postcentral** | **0.3027** | 0.1780 | **0.2952** |
| **lh-G_precentral** | 0.1574 | 0.0539 | 0.1915 |
| **lh-G_precuneus** | 0.0358 | -0.0728 | 0.1495 |
| **lh-G_rectus** | 0.0996 | 0.0335 | 0.0209 |
| **lh-G_subcallosal** | 0.0902 | 0.1587 | -0.1493 |
| **lh-G_temp_sup-G_T_transv** | 0.0928 | 0.1774 | -0.0991 |
| **lh-G_temp_sup-Lateral** | -0.0313 | -0.0364 | 0.0660 |
| **lh-G_temp_sup-Plan_polar** | 0.0113 | 0.0612 | 0.0331 |
| **lh-G_temp_sup-Plan_tempo** | 0.0716 | 0.1113 | 0.0175 |
| **lh-G_temporal_inf** | -0.0260 | -0.0243 | 0.0387 |
| **lh-G_temporal_middle** | -0.1341 | -0.0905 | -0.0042 |
| **lh-Lat_Fis-ant-Horizont** | 0.0469 | -0.0491 | 0.0610 |
| **lh-Lat_Fis-ant-Vertical** | 0.1610 | 0.1226 | 0.1431 |

| | | | |
|---|---|---|---|
| **lh-Lat_Fis-post** | 0.0522 | 0.1219 | -0.1070 |
| **lh-Pole_occipital** | -0.0569 | -0.1347 | 0.0897 |
| **lh-Pole_temporal** | -0.1291 | -0.0909 | -0.1691 |
| **lh-S_calcarine** | 0.1104 | 0.1855 | -0.1190 |
| **lh-S_central** | 0.1844 | 0.1395 | 0.1888 |
| **lh-S_cingul-Marginalis** | -0.0186 | -0.0533 | 0.0877 |
| **lh-S_circular_insula_ant** | 0.0683 | 0.1270 | -0.1403 |
| **lh-S_circular_insula_inf** | -0.0035 | 0.0513 | 0.1384 |
| **lh-S_circular_insula_sup** | 0.0018 | 0.0287 | 0.0493 |
| **lh-S_collat_transv_ant** | -0.0074 | -0.0092 | 0.0933 |
| **lh-S_collat_transv_post** | 0.0843 | 0.0586 | 0.1399 |
| **lh-S_front_inf** | -0.0185 | -0.0651 | 0.2502 |
| **lh-S_front_middle** | 0.0114 | -0.0434 | **0.3667** |
| **lh-S_front_sup** | 0.1565 | 0.0784 | **0.3611** |
| **lh-S_interm_prim-Jensen** | -0.0962 | -0.0796 | -0.0162 |
| **lh-S_intrapariet&P_trans** | 0.0721 | 0.0879 | 0.0486 |
| **lh-S_oc-temp_lat** | -0.0593 | -0.0401 | 0.0746 |
| **lh-S_oc-temp_med&Lingual** | 0.0855 | 0.1097 | 0.0161 |
| **lh-S_oc_middle&Lunatus** | -0.0103 | -0.0078 | 0.0515 |
| **lh-S_oc_sup&transversal** | 0.1353 | 0.1181 | -0.0291 |
| **lh-S_occipital_ant** | -0.0098 | 0.0274 | 0.0358 |
| **lh-S_orbital-H_Shaped** | 0.0703 | -0.0309 | 0.1129 |
| **lh-S_orbital_lateral** | -0.0136 | 0.0377 | -0.0764 |
| **lh-S_orbital_med-olfact** | -0.0727 | -0.1375 | 0.1312 |
| **lh-S_parieto_occipital** | 0.0610 | 0.0562 | -0.0578 |
| **lh-S_pericallosal** | -0.1091 | -0.0351 | -0.1058 |
| **lh-S_postcentral** | 0.1341 | 0.0319 | **0.3347** |
| **lh-S_precentral-inf-part** | 0.1881 | 0.1218 | 0.2517 |
| **lh-S_precentral-sup-part** | 0.1268 | 0.0455 | 0.2775 |
| **lh-S_suborbital** | -0.1288 | 0.0159 | -0.1953 |
| **lh-S_subparietal** | -0.0646 | -0.0484 | 0.0172 |
| **lh-S_temporal_inf** | 0.0126 | 0.0374 | 0.0635 |
| **lh-S_temporal_sup** | -0.0749 | -0.0646 | 0.0177 |
| **lh-S_temporal_transverse** | -0.0545 | 0.0387 | -0.1295 |
| **rh-G&S_cingul-Ant** | -0.0549 | 0.0579 | -0.0652 |
| **rh-G&S_cingul-Mid-Ant** | -0.1847 | -0.1642 | 0.0525 |
| **rh-G&S_cingul-Mid-Post** | -0.2259 | -0.2268 | -0.0911 |
| **rh-G&S_frontomargin** | 0.1479 | 0.0452 | 0.2066 |
| **rh-G&S_occipital_inf** | -0.0863 | -0.0655 | -0.0064 |
| **rh-G&S_paracentral** | 0.2322 | 0.0478 | 0.2640 |
| **rh-G&S_subcentral** | -0.1209 | -0.0993 | -0.0626 |

| | | | |
|---|---|---|---|
| **rh-G&S_transv_frontopol** | 0.2181 | 0.1562 | 0.1858 |
| **rh-G_Ins_lg&S_cent_ins** | -0.0395 | -0.0168 | -0.0005 |
| **rh-G_cingul-Post-dorsal** | -0.0815 | -0.0442 | -0.1065 |
| **rh-G_cingul-Post-ventral** | -0.0107 | 0.0415 | 0.0060 |
| **rh-G_cuneus** | 0.2036 | 0.0539 | 0.2125 |
| **rh-G_front_inf-Opercular** | -0.1621 | -0.0826 | -0.1097 |
| **rh-G_front_inf-Orbital** | -0.0797 | -0.0485 | -0.0637 |
| **rh-G_front_inf-Triangul** | 0.0758 | 0.0537 | 0.0541 |
| **rh-G_front_middle** | 0.1044 | 0.0448 | 0.1799 |
| **rh-G_front_sup** | -0.0366 | -0.1163 | 0.1651 |
| **rh-G_insular_short** | -0.0962 | -0.0546 | -0.0509 |
| **rh-G_oc-temp_lat-fusifor** | 0.0192 | -0.0179 | 0.0926 |
| **rh-G_oc-temp_med-Lingual** | 0.0906 | 0.2130 | -0.1277 |
| **rh-G_oc-temp_med-Parahip** | 0.0858 | 0.0730 | 0.0681 |
| **rh-G_occipital_middle** | 0.0341 | 0.0419 | -0.0845 |
| **rh-G_occipital_sup** | -0.0380 | -0.0897 | 0.0183 |
| **rh-G_orbital** | 0.0753 | 0.0893 | -0.0039 |
| **rh-G_pariet_inf-Angular** | -0.0624 | 0.0598 | -0.1902 |
| **rh-G_pariet_inf-Supramar** | -0.0721 | -0.0368 | -0.0798 |
| **rh-G_parietal_sup** | 0.1795 | 0.0978 | 0.1955 |
| **rh-G_postcentral** | 0.1523 | 0.0274 | 0.1851 |
| **rh-G_precentral** | -0.0528 | -0.1387 | 0.1283 |
| **rh-G_precuneus** | 0.2207 | 0.1774 | 0.1010 |
| **rh-G_rectus** | 0.0700 | -0.0398 | 0.0555 |
| **rh-G_subcallosal** | -0.0868 | -0.0449 | -0.0754 |
| **rh-G_temp_sup-G_T_transv** | -0.0966 | -0.0362 | -0.1228 |
| **rh-G_temp_sup-Lateral** | -0.0367 | -0.0738 | 0.0927 |
| **rh-G_temp_sup-Plan_polar** | 0.0577 | 0.0792 | 0.0998 |
| **rh-G_temp_sup-Plan_tempo** | -0.0848 | -0.0255 | -0.0792 |
| **rh-G_temporal_inf** | -0.0216 | 0.0167 | -0.1030 |
| **rh-G_temporal_middle** | -0.1283 | -0.0796 | -0.0724 |
| **rh-Lat_Fis-ant-Horizont** | -0.0797 | -0.0521 | -0.0716 |
| **rh-Lat_Fis-ant-Vertical** | -0.0288 | -0.0265 | 0.0181 |
| **rh-Lat_Fis-post** | -0.1607 | -0.1453 | -0.0078 |
| **rh-Pole_occipital** | 0.0130 | -0.0396 | 0.1128 |
| **rh-Pole_temporal** | -0.0632 | -0.1114 | 0.0621 |
| **rh-S_calcarine** | 0.1710 | 0.2205 | -0.0850 |
| **rh-S_central** | 0.0652 | 0.0417 | 0.1172 |
| **rh-S_cingul-Marginalis** | 0.1391 | 0.1402 | 0.1510 |
| **rh-S_circular_insula_ant** | -0.1450 | -0.0914 | -0.1651 |

| | | | |
|---|---|---|---|
| **rh-S_circular_insula_inf** | 0.0365 | 0.0900 | -0.0696 |
| **rh-S_circular_insula_sup** | -0.0216 | -0.0368 | 0.0580 |
| **rh-S_collat_transv_ant** | 0.0183 | 0.0027 | -0.0230 |
| **rh-S_collat_transv_post** | 0.1182 | 0.0493 | 0.1308 |
| **rh-S_front_inf** | -0.0198 | -0.0716 | 0.2363 |
| **rh-S_front_middle** | 0.2624 | 0.1831 | **0.3281** |
| **rh-S_front_sup** | 0.2530 | 0.1272 | **0.3443** |
| **rh-S_interm_prim-Jensen** | 0.0881 | 0.0673 | 0.0525 |
| **rh-S_intrapariet&P_trans** | 0.1992 | 0.1458 | 0.2553 |
| **rh-S_oc-temp_lat** | 0.0138 | 0.0319 | -0.0156 |
| **rh-S_oc-temp_med&Lingual** | 0.0910 | 0.0677 | 0.1303 |
| **rh-S_oc_middle&Lunatus** | -0.0215 | -0.0337 | 0.0705 |
| **rh-S_oc_sup&transversal** | -0.0997 | -0.1088 | 0.0121 |
| **rh-S_occipital_ant** | -0.0093 | 0.0390 | -0.0531 |
| **rh-S_orbital-H_Shaped** | 0.0916 | 0.0095 | 0.2054 |
| **rh-S_orbital_lateral** | 0.0180 | -0.0404 | 0.2653 |
| **rh-S_orbital_med-olfact** | -0.1298 | -0.0971 | -0.0152 |
| **rh-S_parieto_occipital** | 0.2387 | 0.1757 | 0.2379 |
| **rh-S_pericallosal** | -0.1621 | -0.1205 | -0.1081 |
| **rh-S_postcentral** | 0.0392 | -0.0360 | **0.3153** |
| **rh-S_precentral-inf-part** | -0.0909 | -0.1162 | 0.1639 |
| **rh-S_precentral-sup-part** | 0.0376 | -0.0508 | **0.3139** |
| **rh-S_suborbital** | 0.0579 | 0.1742 | -0.0994 |
| **rh-S_subparietal** | 0.0510 | 0.1175 | -0.0740 |
| **rh-S_temporal_inf** | -0.0436 | -0.0061 | -0.0945 |
| **rh-S_temporal_sup** | -0.1917 | -0.1603 | -0.1881 |
| **rh-S_temporal_transverse** | 0.0362 | 0.0339 | -0.0053 |

**Table S5**: Pearson r correlation between regional saliency and gray matter, surface area, and cortical thickness. Significant correlations after Benjamini-Hochberg FDR correction are bolded.

| Measure | Group | n subjects | n regions | Mantel r | Mantel p | Mantel p_fdr |
|---|---|---|---|---|---|---|
| Cortical thickness | all | 132 | 148 | 0.22426 | 0.0005 | 0.0009 |
| | low | 72 | 148 | 0.262996 | 0.0005 | 0.0009 |
| | high | 60 | 148 | 0.137902 | 0.004498 | 0.00506 |
| Surface area | all | 132 | 148 | 0.139805 | 0.001999 | 0.00257 |
| | low | 72 | 148 | 0.124813 | 0.001499 | 0.002249 |
| | high | 60 | 148 | 0.108166 | 0.013993 | 0.013993 |
| Gray matter volume | all | 132 | 148 | 0.2136 | 0.0005 | 0.0009 |
| | low | 72 | 148 | 0.191629 | 0.0005 | 0.0009 |
| | high | 60 | 148 | 0.168439 | 0.0005 | 0.0009 |

**Table S6**: Inter-structural-covariance-analysis (ISCA) results between saliency covariation networks (all subjects, low-severity subjects, high-severity subjects) and structural covariance networks (cortical thickness, surface area, gray matter volume). Mantel r, Mantel p, and Mantel p after Benjamini-Hochberg FDR correction are reported.

| τ > 0.1: Region | Correlations | τ > 0.2: Region | Correlations | τ > 0.3: Region | Correlations |
|---|---|---|---|---|---|
| lh_G_front_inf-Opercular | 147 | lh_G_Ins_lg_and_S_cent_ins | 146 | lh_G_and_S_subcentral | 137 |
| lh_G_rectus | 147 | lh_S_circular_insula_inf | 144 | lh_G_and_S_cingul-Ant | 137 |
| lh_G_and_S_cingul-Ant | 147 | lh_S_circular_insula_sup | 144 | lh_G_temp_sup-G_T_transv | 135 |
| lh_G_Ins_lg_and_S_cent_ins | 147 | lh_G_front_inf-Opercular | 143 | lh_G_and_S_cingul-Mid-Ant | 135 |
| lh_G_insular_short | 147 | lh_S_precentral-inf-part | 143 | lh_S_central | 134 |
| lh_S_circular_insula_sup | 147 | lh_G_and_S_cingul-Ant | 143 | lh_S_pericallosal | 134 |
| lh_G_precentral | 146 | lh_G_subcallosal | 143 | rh_G_subcallosal | 134 |
| lh_S_front_middle | 146 | lh_G_and_S_subcentral | 142 | lh_G_precentral | 133 |
| lh_S_front_sup | 146 | lh_G_front_sup | 142 | lh_Lat_Fis-post | 133 |
| lh_S_suborbital | 146 | lh_S_suborbital | 142 | lh_S_temporal_sup | 132 |
| lh_G_postcentral | 146 | lh_G_and_S_cingul-Mid-Ant | 142 | lh_G_pariet_inf-Supramar | 132 |
| lh_S_precentral-inf-part | 146 | lh_G_precentral | 141 | lh_G_and_S_cingul-Mid-Post | 132 |
| lh_S_precentral-sup-part | 146 | lh_G_temp_sup-G_T_transv | 141 | lh_S_circular_insula_inf | 132 |
| lh_G_subcallosal | 146 | lh_S_pericallosal | 141 | rh_G_insular_short | 132 |
| lh_S_circular_insula_inf | 146 | lh_G_and_S_cingul-Mid-Post | 141 | rh_S_orbital-H_Shaped | 132 |
| lh_G_and_S_subcentral | 145 | rh_G_subcallosal | 141 | lh_G_temp_sup-Plan_tempo | 131 |
| lh_G_front_middle | 145 | rh_S_orbital-H_Shaped | 141 | lh_G_postcentral | 131 |
| lh_G_front_sup | 145 | lh_S_central | 140 | lh_S_oc_sup_and_transversal | 131 |
| lh_G_and_S_cingul-Mid-Ant | 145 | lh_S_front_middle | 140 | rh_S_circular_insula_ant | 131 |
| rh_G_and_S_transv_frontopol | 145 | lh_S_front_sup | 140 | lh_S_cingul-Marginalis | 130 |
| lh_S_central | 144 | lh_G_temp_sup-Plan_tempo | 140 | rh_S_circular_insula_sup | 130 |
| lh_G_temporal_middle | 144 | lh_G_postcentral | 140 | lh_S_temporal_transverse | 129 |
| lh_S_pericallosal | 144 | lh_S_precentral-sup-part | 140 | lh_S_subparietal | 129 |
| lh_G_and_S_cingul-Mid-Post | 144 | lh_G_insular_short | 140 | lh_G_occipital_sup | 129 |
| lh_S_circular_insula_ant | 144 | rh_G_orbital | 140 | lh_G_Ins_lg_and_S_cent_ins | 129 |
| rh_G_subcallosal | 144 | lh_Lat_Fis-post | 139 | rh_G_and_S_cingul-Mid-Ant | 129 |
| rh_S_pericallosal | 144 | lh_S_temporal_sup | 139 | rh_Pole_occipital | 129 |
| rh_Pole_temporal | 144 | lh_G_pariet_inf-Supramar | 139 | rh_G_temporal_middle | 129 |
| rh_S_orbital_med-olfact | 144 | rh_S_circular_insula_ant | 139 | rh_G_front_inf-Orbital | 129 |
| rh_G_front_sup | 144 | rh_G_and_S_cingul-Mid-Post | 139 | lh_S_intrapariet_and_P_trans | 128 |
| rh_G_and_S_frontomargin | 144 | rh_G_and_S_cingul-Mid-Ant | 139 | lh_G_occipital_middle | 128 |
| lh_S_front_inf | 143 | rh_S_pericallosal | 139 | lh_S_oc_middle_and_Lunatus | 128 |
| lh_S_orbital_med-olfact | 143 | rh_G_and_S_frontomargin | 139 | lh_G_cingul-Post-dorsal | 128 |
| lh_G_temp_sup-G_T_transv | 143 | lh_S_temporal_transverse | 138 | rh_G_and_S_cingul-Mid-Post | 128 |
| lh_S_temporal_sup | 143 | lh_S_postcentral | 138 | rh_G_and_S_occipital_inf | 128 |
| lh_G_pariet_inf-Supramar | 143 | rh_G_insular_short | 138 | rh_G_temp_sup-Plan_tempo | 128 |
| rh_S_circular_insula_ant | 143 | rh_G_and_S_occipital_inf | 138 | rh_G_and_S_subcentral | 128 |
| rh_G_cingul-Post-dorsal | 143 | rh_G_temp_sup-Plan_polar | 138 | lh_S_front_sup | 127 |
| rh_G_and_S_cingul-Mid-Post | 143 | rh_S_orbital_med-olfact | 138 | lh_G_oc-temp_med-Lingual | 127 |
| rh_G_and_S_cingul-Ant | 143 | rh_S_front_middle | 138 | lh_G_pariet_inf-Angular | 127 |
| rh_G_temp_sup-Plan_polar | 143 | lh_S_subparietal | 137 | rh_G_Ins_lg_and_S_cent_ins | 127 |
| rh_S_orbital-H_Shaped | 143 | lh_G_and_S_occipital_inf | 137 | rh_S_pericallosal | 127 |
| rh_G_orbital | 143 | lh_G_occipital_middle | 137 | rh_Lat_Fis-post | 127 |
| lh_Lat_Fis-post | 142 | lh_G_occipital_sup | 137 | rh_G_orbital | 127 |
| lh_S_temporal_transverse | 142 | lh_S_occipital_ant | 137 | lh_G_parietal_sup | 126 |
| lh_S_postcentral | 142 | lh_G_cingul-Post-dorsal | 137 | lh_S_parieto_occipital | 126 |
| lh_S_subparietal | 142 | rh_S_circular_insula_sup | 137 | lh_G_and_S_occipital_inf | 126 |
| lh_G_occipital_sup | 142 | rh_G_Ins_lg_and_S_cent_ins | 137 | lh_S_calcarine | 126 |
| lh_G_cingul-Post-dorsal | 142 | rh_Lat_Fis-ant-Horizont | 137 | rh_S_circular_insula_inf | 126 |
| lh_S_cingul-Marginalis | 142 | lh_S_intrapariet_and_P_trans | 136 | rh_S_cingul-Marginalis | 126 |

| $\tau > 0.4$: Region | Correlations | $\tau > 0.5$: Region | Correlations | $\tau > 0.6$: Region | Correlations |
|---|---|---|---|---|---|
| lh_G_and_S_subcentral | 125 | rh_Lat_Fis-post | 90 | lh_Lat_Fis-post | 32 |
| lh_G_pariet_inf-Supramar | 123 | rh_S_circular_insula_sup | 87 | rh_Lat_Fis-post | 28 |
| rh_S_circular_insula_sup | 121 | lh_Lat_Fis-post | 84 | lh_G_oc-temp_med-Lingual | 26 |
| lh_G_and_S_cingul-Mid-Post | 119 | rh_G_insular_short | 80 | rh_S_circular_insula_sup | 26 |
| rh_Lat_Fis-post | 118 | rh_G_Ins_lg_and_S_cent_ins | 80 | lh_S_calcarine | 25 |
| lh_Lat_Fis-post | 117 | lh_G_oc-temp_med-Lingual | 77 | lh_G_parietal_sup | 24 |
| lh_S_pericallosal | 117 | lh_G_pariet_inf-Supramar | 77 | lh_S_intrapariet_and_P_trans | 24 |
| lh_G_occipital_sup | 117 | lh_S_calcarine | 74 | rh_G_insular_short | 23 |
| rh_G_insular_short | 117 | lh_G_and_S_cingul-Mid-Post | 74 | rh_G_Ins_lg_and_S_cent_ins | 23 |
| lh_S_central | 116 | lh_S_cingul-Marginalis | 74 | lh_G_pariet_inf-Supramar | 22 |
| lh_G_temp_sup-G_T_transv | 115 | rh_S_precentral-inf-part | 73 | lh_G_occipital_middle | 21 |
| lh_S_subparietal | 114 | lh_G_occipital_sup | 72 | lh_S_cingul-Marginalis | 21 |
| rh_S_front_middle | 114 | rh_S_front_middle | 72 | rh_S_central | 21 |
| lh_S_temporal_sup | 113 | lh_S_temporal_transverse | 71 | lh_S_central | 20 |
| lh_G_pariet_inf-Angular | 113 | lh_G_cuneus | 71 | lh_G_temp_sup-G_T_transv | 20 |
| lh_S_cingul-Marginalis | 113 | rh_Pole_occipital | 71 | lh_S_temporal_sup | 20 |
| rh_S_circular_insula_ant | 112 | lh_G_temp_sup-G_T_transv | 70 | lh_G_cuneus | 20 |
| lh_S_oc_sup_and_transversal | 111 | lh_S_temporal_sup | 70 | lh_S_oc-temp_med_and_Lingual | 20 |
| lh_G_and_S_cingul-Mid-Ant | 111 | lh_G_occipital_middle | 70 | lh_S_parieto_occipital | 19 |
| lh_G_occipital_middle | 110 | rh_S_circular_insula_inf | 70 | rh_S_temporal_sup | 19 |
| rh_S_cingul-Marginalis | 110 | lh_G_temp_sup-Plan_tempo | 68 | lh_S_temporal_transverse | 18 |
| rh_S_precentral-inf-part | 110 | lh_S_subparietal | 68 | lh_S_precentral-sup-part | 18 |
| lh_S_parieto_occipital | 109 | rh_S_cingul-Marginalis | 68 | lh_G_temp_sup-Plan_tempo | 17 |
| lh_S_oc_middle_and_Lunatus | 109 | lh_S_intrapariet_and_P_trans | 66 | lh_G_pariet_inf-Angular | 17 |
| rh_Lat_Fis-ant-Horizont | 109 | lh_S_oc-temp_med_and_Lingual | 66 | rh_S_circular_insula_inf | 17 |
| rh_G_and_S_subcentral | 109 | rh_S_front_inf | 66 | rh_S_postcentral | 17 |
| rh_G_front_inf-Opercular | 108 | lh_G_pariet_inf-Angular | 65 | rh_S_intrapariet_and_P_trans | 17 |
| lh_S_interm_prim-Jensen | 107 | lh_S_parieto_occipital | 65 | rh_S_front_sup | 17 |
| rh_G_subcallosal | 107 | lh_G_and_S_subcentral | 64 | lh_G_and_S_subcentral | 16 |
| rh_G_and_S_cingul-Mid-Ant | 107 | lh_S_pericallosal | 64 | lh_G_and_S_cingul-Mid-Post | 16 |
| rh_G_precuneus | 107 | rh_G_front_inf-Opercular | 64 | rh_S_precentral-inf-part | 16 |
| rh_G_temp_sup-Plan_tempo | 107 | lh_S_central | 63 | rh_S_front_inf | 16 |
| lh_S_intrapariet_and_P_trans | 106 | rh_G_subcallosal | 63 | rh_G_front_inf-Opercular | 16 |
| rh_Pole_occipital | 106 | lh_G_parietal_sup | 62 | lh_G_postcentral | 15 |
| rh_S_temporal_sup | 106 | lh_S_oc_sup_and_transversal | 62 | lh_S_oc_sup_and_transversal | 15 |
| rh_S_front_inf | 106 | rh_G_and_S_cingul-Mid-Ant | 62 | lh_S_postcentral | 14 |
| lh_G_temp_sup-Plan_tempo | 105 | rh_G_precuneus | 62 | lh_S_precentral-inf-part | 13 |
| lh_Pole_occipital | 105 | rh_S_temporal_sup | 62 | rh_G_parietal_sup | 13 |
| lh_S_calcarine | 105 | lh_S_interm_prim-Jensen | 61 | rh_S_precentral-sup-part | 12 |
| rh_G_Ins_lg_and_S_cent_ins | 104 | lh_G_and_S_occipital_inf | 61 | rh_G_and_S_subcentral | 12 |
| rh_G_temporal_middle | 104 | rh_G_and_S_cingul-Mid-Post | 61 | lh_G_precentral | 11 |
| rh_G_and_S_cingul-Mid-Post | 103 | rh_G_temporal_middle | 59 | lh_G_and_S_occipital_inf | 11 |
| rh_G_front_inf-Orbital | 103 | lh_Pole_occipital | 58 | lh_G_and_S_cingul-Mid-Ant | 11 |
| lh_G_oc-temp_med-Lingual | 102 | lh_G_precuneus | 54 | lh_S_circular_insula_sup | 11 |
| lh_S_circular_insula_inf | 102 | rh_S_parieto_occipital | 54 | rh_G_subcallosal | 11 |
| rh_G_and_S_occipital_inf | 102 | rh_G_pariet_inf-Supramar | 54 | rh_G_postcentral | 11 |
| rh_G_pariet_inf-Supramar | 102 | lh_S_oc-temp_lat | 53 | rh_G_pariet_inf-Supramar | 11 |
| lh_S_temporal_transverse | 101 | rh_G_temp_sup-G_T_transv | 53 | lh_S_front_sup | 10 |
| rh_S_circular_insula_inf | 101 | lh_G_oc-temp_lat-fusifor | 52 | lh_G_oc-temp_lat-fusifor | 10 |
| rh_S_orbital-H_Shaped | 101 | rh_S_circular_insula_ant | 52 | lh_S_interm_prim-Jensen | 10 |

| τ > 0.7: Region | Correlations |
|---|---|
| lh_G_precentral | 5 |
| lh_G_temp_sup-G_T_transv | 5 |
| lh_S_precentral-sup-part | 5 |
| rh_S_precentral-sup-part | 5 |
| rh_S_central | 5 |
| lh_S_central | 4 |
| lh_G_postcentral | 4 |
| lh_S_postcentral | 4 |
| lh_S_precentral-inf-part | 4 |
| lh_S_circular_insula_sup | 4 |
| rh_S_postcentral | 4 |
| rh_S_intrapariet_and_P_trans | 4 |
| lh_S_front_sup | 3 |
| lh_G_oc-temp_med-Lingual | 3 |
| lh_S_temporal_sup | 3 |
| lh_S_temporal_transverse | 3 |
| lh_S_intrapariet_and_P_trans | 3 |
| lh_S_circular_insula_ant | 3 |
| rh_G_parietal_sup | 3 |
| rh_G_and_S_subcentral | 3 |
| lh_G_front_inf-Opercular | 2 |
| lh_Lat_Fis-post | 2 |
| lh_G_temp_sup-Plan_tempo | 2 |
| lh_G_pariet_inf-Angular | 2 |
| lh_G_insular_short | 2 |
| lh_S_cingul-Marginalis | 2 |
| lh_S_circular_insula_inf | 2 |
| rh_S_circular_insula_sup | 2 |
| rh_S_circular_insula_ant | 2 |
| rh_G_temp_sup-Lateral | 2 |
| rh_S_orbital-H_Shaped | 2 |
| rh_S_front_sup | 2 |
| rh_G_precentral | 2 |
| lh_G_and_S_subcentral | 1 |
| lh_G_front_inf-Orbital | 1 |
| lh_Lat_Fis-ant-Horizont | 1 |
| lh_S_front_inf | 1 |
| lh_S_front_middle | 1 |
| lh_S_orbital-H_Shaped | 1 |
| lh_S_suborbital | 1 |
| lh_G_oc-temp_lat-fusifor | 1 |
| lh_G_temporal_inf | 1 |
| lh_S_temporal_inf | 1 |
| lh_G_pariet_inf-Supramar | 1 |
| lh_S_subparietal | 1 |
| lh_G_cuneus | 1 |
| lh_G_occipital_middle | 1 |
| lh_S_calcarine | 1 |
| lh_S_oc-temp_med_and_Lingual | 1 |
| lh_G_and_S_cingul-Ant | 1 |

**Table S7:** Ordered list of top 50 cortical regions ranked by their number of significant saliency correlations with other brain regions passing each of 7 correlation coefficient thresholds τ = 0.1, 0.2,…, 0.7.

| Subscore | Hemisphere | Lobe | Area | n | tau | p |
|---|---|---|---|---|---|---|
| **Cognition** | lh | frontal | G_orbital | 132 | 0.1268 | 0.0348 |
| | lh | temporal | G_temp_sup-Plan_polar | 132 | 0.1216 | 0.0429 |
| | lh | parietal | S_interm_prim-Jensen | 132 | -0.1649 | 0.006 |
| | rh | occipital | S_oc-temp_med_and_Lingual | 132 | -0.1237 | 0.0394 |
| **Awareness** | lh | parietal | S_interm_prim-Jensen | 132 | -0.1379 | 0.0206 |
| **Communication** | lh | temporal | G_temp_sup-Plan_tempo | 132 | -0.1299 | 0.0295 |
| | lh | parietal | S_interm_prim-Jensen | 132 | -0.186 | 0.0018 |
| | lh | parietal | G_pariet_inf-Supramar | 132 | -0.1324 | 0.0265 |
| | lh | parietal | S_pericallosal | 132 | -0.1266 | 0.0339 |
| | lh | occipital | S_oc-temp_lat | 132 | -0.1186 | 0.0469 |
| | rh | temporal | G_temporal_inf | 132 | -0.1284 | 0.0314 |
| | rh | temporal | S_collat_transv_ant | 132 | -0.1181 | 0.0478 |
| | rh | parietal | G_precuneus | 132 | -0.1296 | 0.0298 |
| | rh | parietal | S_pericallosal | 132 | -0.1292 | 0.0304 |
| | rh | parietal | S_subparietal | 132 | -0.1197 | 0.0448 |
| | rh | occipital | S_oc-temp_med_and_Lingual | 132 | -0.1371 | 0.0215 |
| | rh | occipital | G_cuneus | 132 | -0.1331 | 0.0257 |
| | rh | occipital | S_calcarine | 132 | -0.1301 | 0.0293 |
| | rh | occipital | S_oc-temp_lat | 132 | -0.1237 | 0.0381 |
| | rh | limbic | G_cingul-Post-ventral | 132 | -0.1301 | 0.0293 |
| **Mannerism** | lh | temporal | S_collat_transv_post | 132 | -0.1229 | 0.0407 |
| | lh | parietal | S_interm_prim-Jensen | 132 | -0.1661 | 0.0057 |
| | rh | temporal | S_collat_transv_ant | 132 | -0.1201 | 0.0456 |
| | rh | parietal | S_subparietal | 132 | -0.1355 | 0.0241 |
| | rh | occipital | S_oc-temp_med_and_Lingual | 132 | -0.1213 | 0.0435 |
| | rh | occipital | S_calcarine | 132 | -0.118 | 0.0496 |
| **Motivation** | lh | temporal | Pole_temporal | 132 | 0.128 | 0.0333 |
| | lh | temporal | G_temp_sup-Plan_tempo | 132 | -0.1252 | 0.0374 |
| | lh | temporal | G_temp_sup-Plan_polar | 132 | 0.1235 | 0.04 |
| | lh | parietal | S_interm_prim-Jensen | 132 | -0.1795 | 0.0028 |
| | lh | parietal | G_pariet_inf-Supramar | 132 | -0.1185 | 0.0488 |
| | rh | temporal | S_collat_transv_ant | 132 | -0.1276 | 0.034 |
| | rh | occipital | S_oc-temp_med_and_Lingual | 132 | -0.124 | 0.0393 |

**Table S8**: Correlations between regional saliency and SRS subscores (awareness, cognition, communication, mannerism, motivation) by hemisphere and lobe. Correlation did not survive Benjamini-Hochberg FDR correction and are reported as nominal p values.

### Materials and Methods

*Data Acquisition*

We used sMRI sequences from the Autism Brain Imaging Data Exchange II (ABIDE-II), which is a collection of 1114 subjects (521 diagnosed with ASD and 593 control) gathered from 19 institutions. ABIDE-II consists of male and female subjects aging from 5 years old to 64 years old. Compared to its previous version ABIDE I, ABIDE-II has a greater phenotypic characterization through including additional measures such as different ASD assessment and phenotypes related to ASD research (Di Martino et al. 2017).

We only considered sequences from subjects that were assessed with the Social Response Scale 2 (SRS-2), which is an assessment for quantifying the severity of five ASD related phenotypes (Awareness, Cognition, Communication, Motivation, and Mannerisms). This assessment can be administered to subjects ranging from ages 2 years and 5 months to 18 years old and by gender. By quantifying the severity of these phenotypes, it is possible to compare the severity of ASD in different subjects. Each phenotype is scored by summing a questionnaire with a 4-point Likert-scale, constituting the raw score for each respective phenotype. The SRS-2 Total T-score is generated by scaling the raw score for each phenotype and combining them together (Bruni 2014).

We considered subjects' data from the KKI, NYU-1, OHSU, SDSU, and TCD sites (only subjects in these sites were assessed with SRS Total T-Score) from the ABIDE-II dataset and subsequently uploaded to the Boston University Shared Computing Cluster (BU SCC). The total number of the subjects with SRS-T Total Scores we accessed from the sites totaled to 420 subjects. Since ASD is more prevalent in the male population than in the female population with a ratio of 3:1 (Loomes, Hull, and Mandy 2017), only male subjects were used in this study. Additionally, subjects with the age 12 years or younger were considered to be in the age range of 6-12 years old. For AI supervised training subjects were labeled based on their SRS Total T-Score; low severity subjects were labeled with an SRS Total T-Score of 45 or less and high severity subjects were labeled with an SRS Total T-Score of 70 or higher (Bruni 2014). These score criteria yielded 132 subjects for our AI training and analyses (Table S1).

*Preprocessing Methods*

We preprocessed MRI data to reduce within and across subject groups (low vs. high severity) variabilities unrelated to pathology, in order to minimize confounding neural network performance for the classification of ASD severity (Table S1-S2). We utilized FreeSurfer (Fischl 2012) and FSL (Smith et al. 2004) to preprocess the MRI volumes. We used Recon-all, a preprocessing pipeline that performs the Freesurfer cortical reconstruction process. These preprocessing steps included:

*Motion Correction*

Human motion that may have been recorded during scanning procedures was corrected for by constructing the mean of multiple scans. Additionally, subject volumes were transformed from their unique native space to anatomical space (1 mm voxels in a 256╳256╳256 space) (Collins et al. 1994).

*Intensity Normalization*

Four iterations of non-parametric non-uniform intensity normalization was performed to reduce fluctuations in intensity through the nu_correct algorithm in recon-all. We used the mri_normalize algorithm to normalize the intensity value so that the white matter peak normalized to a value of 110 (Dale, Fischl, and Sereno 1999).

*Talairach Transformation*

We used MINC program mritotal to calculate affine transformation for the original volume to be aligned with the MNI305 atlas. The subject MRI volume was aligned to the MNI305 template (at 1 mm resolution) in anatomical space. This allowed similar regions in each subject brain to share similar voxel regions within the template (Collins et al. 1994).

*Smoothing*

A multidimensional (3D) gaussian filter was applied to smooth MRI volumes through applying the standard deviation of the gaussian kernel $\sigma$, which can be computed from the width at half maximum *FWHM* with this equation:

$$FWHM = \sqrt{8ln2}\,\sigma,$$

which can be rearranged to solve for $\sigma$:

$$\sigma = \frac{FWHM}{\sqrt{8\ln 2}}.$$

A *FWHM* value of 3 times the voxel size was suggested (Mikl et al. 2008), so a *FWHM* value of 3 ($\sigma = 1.2739$) was used in this study as the MRIs have voxel size of 1 mm.

*Neural Network Architecture*

A pretrained MedicalNet 3D ResNet-50 backbone (trained on 23 medical imaging datasets) (He et al. 2015, Chen et al. 2019) was used as a feature extractor with all backbone weights frozen, with a trainable classification head to predict a binary high-/low-severity prediction from the resulting feature vector. This architecture was used as it reduced the number of parameters learned from the feature vectors, reducing the potential to overfit compared to a 3D CNN trained from scratch.

*Training*

The classification head was trained for 1000 epochs (minimum epochs: 8 epochs, patience: 6 epochs) using Adam optimizer, with a learning rate of $3\times10^{-5}$ and batch size 4. Prior to training, possible confounds including age (Mann-Whitney U), estimated total intracranial volume (t-test),

IQ (t-test), Euler number (Mann-Whitney U), and site composition ($\chi^2$) were compared between high- and low-severity groups with Benjamini-Hochberg FDR correction (Table S3). Five-fold leave-one-site-out cross-validation and adaptive batch normalization (AdaBN) were used to address site imbalances in high-/low-severity distribution across sites (Table S4). In addition, the model was validated by label-permutation test over 1000 permutations, 1000 seed-stability runs (varying data split and initialization of the trainable head), and Euler decoding from the 3D CNN feature vectors. The trainable heads with the lowest validation loss per fold were used for the analyses listed below.

*Saliency Maps*

Gradient-based saliency maps were computed per subject using Integrated Gradients (Sundararajan et al. 2017). Integrated Gradients attribute the model prediction to each voxel in the sMRI through a path integral from a baseline image (empty black volume) to the actual input. This method was chosen over gradient-class-activation approaches such as GradCAM, as they require upsampling from low-resolution deep-layer feature maps in 3D CNNs. Per subject saliency maps were computed using the trained 3D CNN's leave-one-site-out fold that the subject was the test set of, which was determined by the lowest validation loss. Saliency maps were kept in their signed form to preserve the directionality of each voxel's contribution to severity prediction.

Saliency maps were validated through the following: a) model randomization test to compare saliency from the trained model to saliency derived from a fully randomized model at the regional level, b) saliency seed stability to compare whether regional saliency was similar across runs and folds (10 subjects across 100 seed/data-split pairs and Jaccard index respectively), and c) correlation of regional saliency with FreeSurfer's cortical thickness, gray matter volume, and surface area across subjects (Benjamini-Hochberg FDR corrected) (see Table S5).

<u>Average Saliency and Analysis</u>

Masks of 148 cortical regions (74 per hemisphere, after leaving out lh/rh medial-wall) in the Destrieux atlas (150 regions) and 20 total regions (10 per hemisphere) in the FreeSurfer subcortical atlas (ASEG) were generated through FreeSurfer's mri_binarize and mri_mask commands. These masks were aligned to the saliency maps, and the average saliency for each region was calculated by dividing the sum of the regional saliency by the area of the mask. After average saliencies for all regions in all subjects were obtained, the average saliency per region and severity (low v. high) were obtained. Average left hemisphere saliencies in low severity subjects, average left hemisphere saliencies in high severity subjects, average right hemisphere saliencies in low severity subjects, and average right hemisphere saliencies in high severity subjects were compared to determine cortical and network patterns.

Additionally, rank-based Kendall Tau correlation of regional saliency was computed for all regional pairs across subjects (corrected by Benjamini-Hochberg FDR). This allowed us to find potential regions that displayed saliencies at a similar frequency. Correlations ($p\_FDR < 0.05$) were visualized using the circular_layout function in the MNE-Python library, with the strength of the correlations determined by the $\tau$ values. The resulting saliency covariation network was plotted on a spring-cluster plot where nodes represented regions, node size was correlated with the amount of significant correlation with other regions, and length between nodes was inversely correlated with the $\tau$ correlation between the nodes.

Inter-structural-covariance-analysis (ISCA) was conducted to determine whether the saliency covariation network reflected morphological organization. The saliency covariation matrix was compared to structural networks built from cortical thickness, gray matter volume, and surface area by Mantel test (2000 permutations) (Table S6).

To understand which functional networks were affected across ASD severity, we thresholded the saliency covariation network at 0.1 intervals from $\tau$ values = 0.1 - 0.7. At each interval, we assigned the top 50 cortical regions, based on their significant correlations with other cortical regions, to the functional networks (default mode network, dorsal/ventral attention network, frontoparietal network, language network, limbic/emotion network, salience network, sensorimotor network, and social network) that they were involved in (Table S7). We then averaged the number of times each functional network showed up in the top 50 regions across $\tau$ thresholds.

Cortical regions with saliencies that were significantly correlated with SRS subscores were obtained by conducting rank-based correlation between each cortical region saliency and the five subscores (awareness, cognition, communication, mannerism, motivation). Results were filtered to only include results with a *p*-value less than 0.05 (none survived Benjamini-Hochberg FDR correction, results reported as nominal) (Table S8).

Furthermore, we graphed scatterplots of the average and standard deviation regional saliency of the left hemisphere low severity subjects, left hemisphere high severity subjects, right hemisphere low severity subjects, and right hemisphere high severity subjects to compare the consistency of saliency across the different groups. Each point in the scatterplot represents a region and is organized by lobe to determine any lobular biases the saliency might have (Fig. 7).
